# Enteroviral 2C protein is an RNA-stimulated ATPase and uses a two-step mechanism for binding to RNA and ATP

**DOI:** 10.1101/2022.09.28.509975

**Authors:** Calvin Yeager, Griffin Carter, David W Gohara, Neela H Yennawar, Eric J Enemark, Jamie J Arnold, Craig E Cameron

**Affiliations:** Department of Microbiology & Immunology, University of North Carolina at Chapel Hill, Chapel Hill, NC 27599, USA; Department of Biochemistry and Molecular Biology, St. Louis University, St. Louis, MO 63104, USA; The Huck Institutes of the Life Sciences, The Pennsylvania State University, University Park, PA 16802, USA; Department of Biochemistry and Molecular Biology, University of Arkansas for Medical Sciences, Little Rock, AR 72205, USA

**Keywords:** enterovirus, poliovirus, 2C, ATPase, helicase, AAA+, SF3

## Abstract

The enteroviral 2C protein is a therapeutic target, but the absence of a mechanistic framework for this enzyme limits our understanding of inhibitor mechanisms. Here we use poliovirus 2C and a derivative thereof to elucidate the first biochemical mechanism for this enzyme and confirm the applicability of this mechanism to other members of the enterovirus genus. Our biochemical data are consistent with a dimer forming in solution, binding to RNA, which stimulates ATPase activity by increasing the rate of hydrolysis without impacting affinity for ATP substantially. Both RNA and DNA bind to the same or overlapping site on 2C, driven by the phosphodiester backbone, but only RNA stimulates ATP hydrolysis. We propose that RNA binds to 2C driven by the backbone, with reorientation of the ribose hydroxyls occurring in a second step to form the catalytically competent state. 2C also uses a two-step mechanism for binding to ATP. Initial binding is driven by the α and β phosphates of ATP. In the second step, the adenine base and other substituents of ATP are used to organize the active site for catalysis. These studies provide the first biochemical description of determinants driving specificity and catalytic efficiency of a picornaviral 2C ATPase.

## INTRODUCTION

The Enterovirus genus of the Picornavirus family is comprised of 15 species (1,2). Seven of these species contain viruses capable of infecting humans: enteroviruses (EVs) A-D and rhinoviruses A-C. The type species of this genus is EV-C, with poliovirus (PV), the causative agent of poliomyelitis, representing the prototype virus. The decades-long global effort to eradicate PV has yet to be completely successful. Importantly, this effort has had no impact on minimizing the rising global incidence of other EVs. EVs with a tropism for the respiratory system and capability of causing acute flaccid myelitis, for example EV-A71 and EV-D68, have been on the rise (3,4). Coxsackievirus B3 (CVB3), a species B EV, continues to be a major cause of viral myocarditis and dilated cardiomyopathy (5,6). If the current severe acute respiratory syndrome coronavirus 2 (SARS2) pandemic has taught us anything, it is that we should not eliminate any circulating virus species from the list of potential agents of a future pandemic. As a result, there has been a recent call for studies of non-poliovirus enteroviruses to enable development of countermeasures (7). Multiple EVs are also featured prominently on the list of viruses of pandemic potential targeted by the recently formed Antiviral Drug Discovery Centers for Pathogens of Pandemic Concern (8).

The enteroviral 2C protein represents an attractive target for development of pan-enteroviral therapeutics. The sequence of this protein is as conserved as the viral RNA-dependent RNA polymerase (RdRp) (9). The 2C protein has roles in genome replication and viral morphogenesis (10–18). Sub-optimal activity of 2C protein during morphogenesis causes defects to entry and/or uncoating (14). Numerous classes of small molecules targeting 2C have been discovered (15,19–21). How 2C specifically contributes to genome replication and morphogenesis is unknown.

2C is a member of the helicase superfamily 3 (SF3), a family of small viral hexameric helicases with the ATPase active site belonging to the AAA+ (ATPases Associated with various cellular Activities) superfamily (22). Inclusion of 2C in this family makes it likely that 2C would fulfill some type of helicase function, but neither widespread demonstration nor a specific biological target of helicase activity has yet been identified. One assumption has been that the ATPase-fueled helicase activity might function during genome replication to assist the RdRp with secondary structure and/or to facilitate reutilization of negative-strand RNA templates by preventing formation of, or resolving, dsRNA. Similarly, the helicase and/or translocase activity might facilitate encapsidation of viral RNA during morphogenesis. Again, the biological, biochemical, or biophysical evidence to support these assumptions is sparse.

There is indisputable evidence that 2C is an ATPase and that this activity is essential during early and late steps of the viral lifecycle (23–25). Evidence for helicase activity has only been demonstrated by a single group (26). To our knowledge, no one has ever reported translocase activity. Most efforts over the past few decades have focused on assembly and documentation of hexamers (27–29), a feat that has yet to be achieved with high efficiency for any picornavirus. A major complication of the vast majority of biochemical and biophysical studies of 2C protein is the use of recombinant proteins containing large amino-terminal fusions to help overcome insolubility caused by the amphipathic amino terminus of 2C (17,28).

We have achieved expression of full-length PV 2C and truncated derivatives thereof. We have established a biochemical framework to guide characterization of the mechanism of the 2C-mediated ATPase activity and elucidation of the mechanism(s) of action of inhibition. Prominent features of this mechanism include formation of a 2C dimer in the absence of RNA, ATP, and other viral factors. Binding of RNA and ATP to 2C occurs using a two-step process, which suggests specific interactions between 2C, 2’-hydroxyls of RNA, and both the base and ribose of ATP. These key principles revealed by studying PV 2C protein extend to EV-A71, CVB3, and EV-D68.

## MATERIALS AND METHODS

### Reagents

RNA oligonucleotides (listed in **Table S1**) were from Horizon Discovery Ltd. (Dharmacon); DNA oligonucleotides were from Integrated DNA Technologies, Inc.; radiolabeled ATP and derivatives, [α^32^P]-ATP (3,000 Ci/mmol), [α^32^P]-2’-dATP (3,000 Ci/mmol), and [γ-^32^P]-ATP (7000 Ci/mmol), were from Perkin Elmer; restriction enzymes were from New England Biolabs; Phusion DNA polymerase and T4 polynucleotide kinase were from ThermoFisher; nucleoside 5’-triphosphates (ultrapure solutions) were from Cytiva; remdesivir triphosphate and 2’-C-methyl-ATP was provided by Gilead Sciences; β,γ-methylene-ATP and AMP-PNP were from MilliporeSigma; All other chemicals and reagents were from VWR, MilliporeSigma, or Fisher Scientific at the highest grade.

### Construction of 2C bacterial expression plasmids

2C proteins produced in this study were expressed and purified as SUMO-fusions. The pSUMO system allows for production of SUMO fusion proteins containing an amino-terminal affinity tag fused to SUMO that can be purified by affinity chromatography and subsequently processed by the SUMO protease, Ulp1 (30). Briefly, the WT 2C gene was PCR amplified from the PV cDNA, pMO-3 plasmid, using primers listed in **Table S2**. We utilized BsaI and XhoI restriction sites for subcloning of DNA fragments using T4 DNA ligase into the pSUMO expression plasmid and confirmed the final sequence, pSUMO-PV-WT-2C, by Sanger sequencing at the Penn State Genomics Core Facility in University Park, PA, or Genewiz at Wake County, NC. The N-terminal derivatives, NΔ39 2C and NΔ115 2C, were cloned using a similar procedure as the WT 2C gene. Site-directed mutants, e.g. pSUMO-PV-2C-D177A, were produced by Quikchange mutagenesis. The genes for 2C from CVB3 (Genbank: U57056.1), EV-A71 (Genbank: MG756724.1), and EV-D68 (Genbank: KM881710.2) were PCR amplified from viral cDNA templates (31–33) and cloned into pSUMO expression plasmid by IN-FUSION (Takara Bio). The final constructs were confirmed by sanger sequencing performed by Genewiz.

### Modeling of PV 2C WT

The WT 2C model was modeled by using ATPγS-bound VAT complex (PDB: 5G4G) and generated using Phyre2 in the Intensive mode (34). The model was then minimized using Schrödinger Maestro, by Schrödringer, Inc.

### Expression and purification of PV 2C WT, PV 2C NΔ39, and other NΔ39 constructs

2*C* protein was expressed in Rosetta (DE3) competent cells (MilliporeSigma). Cells were grown in NZCYM media, pH 7.6, to an optical density (OD_600_) of 1.0 and induced by autoinduction for 48 hours at 15 °C (35). The induction and subsequent purity of 2C fractions was evaluated by Coomassie-stained SDS-PAGE (15 %) gels. Cells were harvested by centrifugation at 5,400 x *g* for 10 min at 4 °C and washed with a buffer containing 10 mM Tris and 1 mM EDTA at pH 8.0 and then re-centrifuged. The cell pellet was resuspended and homogenized in Buffer A [50 mM 3-(Cyclohexylamino)-1-propanesulfonic acid (CAPS) at pH 10.0, 10% (v/v) glycerol, 500 mM NaCl, 5 mM imidazole, 5 mM β-mercaptoethanol (BME), 1 mM EDTA, 1.4 μg/mL pepstatin A, 1.0 μg/mL leupeptin] by Dounce homogenizer at a ratio of 5 mL Buffer A per 1-gram wet cell pellet. Lysis was achieved by passing the suspension through a French press twice at a pressure of 15,000 psi. Phenylmethylsulfonyl fluoride (PMSF) and nonyl phenoxypolyethoxylethanol (NP-40) was added to the cell lysate at a final concentration of 1 mM and 0.1%, respectively. While stirring, polyethylenimine (PEI) was added gradually to a final concentration of 0.25% (w/v) at 4 °C to precipitate any contaminating nucleic acids. The PEI-containing lysate was clarified by centrifugation at 75,000 x *g* for 30 min at 4 °C. Ammonium sulfate was gradually added to the clarified lysate at 4 °C to 40% saturation while stirring to precipitate the protein and remove the PEI. The solution was centrifuged at 75,000 x *g* at 4 °C for 30 min to collect the ammonium sulfate-protein pellet. From this point onward, buffers contained detergent only for purification of PV 2C WT and mutant derivatives. The pellet was resuspended in Buffer B [20 mM HEPES at pH 7.5, 20% (v/v) glycerol, 500 mM NaCl, 5 mM BME, 0.1% (v/v) NP-40] containing 5 mM imidazole. Using a peristaltic pump, a nickel-nitrilotriacetic (Ni-NTA) resin (Thermo Fisher Scientific Inc.) was equilibrated with 5 column volumes (CVs) of Buffer B containing 5 mM imidazole at 1 mL/min. The equilibrated Ni-NTA resin was then mixed with the resuspended protein pellet for 30 min while stirring, and subsequently repacked. To remove contaminants, the loaded resin was washed with 5 CVs Buffer B containing 5 mM imidazole and 4 CVs Buffer B containing 50 mM imidazole. 2C was eluted into multiple fractions with Buffer B containing 500 mM imidazole. 2C protein fractions were pooled, treated with 1 μg ubiquitin-like-specific-protease 1 (ULP-1) per 1 mg protein of interest, and dialyzed against Buffer C [20 mM HEPES at pH 7.5, 20% (v/v) glycerol, 5 mM BME, 50 mM NaCl, 0.1% (v/v) NP40] overnight at 4 °C by using a 12-14,000 MWCO dialysis membrane (Spectrum Laboratories). The dialyzed 2C protein was centrifuged at 75,000 x *g* at 4 °C for 30 min to remove insoluble protein. A Q-column (ThermoFisher Scientific Inc.) was equilibrated with 5 CVs Buffer C. PV 2C protein was passed thru the Q-column as flow-through while the cleaved-SUMO protein bound to the Q-column. A phosphocellulose column was prepared as previously described (36) and equilibrated with buffer C. PV 2C was loaded onto the phosphocellulose column with buffer D [20 mM HEPES at pH 7.5, 20% (v/v) glycerol, 5 mM BME, 1.5% (v/v) octyl β-D-glucopyranoside] containing 50 mM NaCl until A_280_ values from the NP-40 were reduced to < 0.1. PV 2C was eluted from the phosphocellulose column with buffer D containing 500 mM NaCl. The protein concentration was determined by measuring the absorbance at 280 nm (PV 2C WT: *ε_max_* = 32,430 M^-1^·cm^-1^·PV 2C NΔ39: *ε_max_* = 15,930 M^-1^·cm^-1^·CVB3 2C NΔ39: *ε_max_* = 14,900 M^-1^·cm^-1^; EVA71 2C NΔ39: *ε_max_* = 14,900 M^-1^·cm^-1^; EVD68 2C NΔ39: *ε_max_* = 20,860 M^-1^·cm^-1^) with NanoDrop One (Thermo Fisher Scientific Inc.). The ionic strength of the dialyzed protein solution was confirmed with a conductivity meter. Agarose gel electrophoresis was run to check for any contaminating nucleic acids. Flash frozen aliquots were stored at −80 °C until ready to use. The ProtParam tool (ExPASy Bioinformatics Resource Portal) was used to calculate the physical/chemical parameters of the protein including extinction coefficient. Representative purified proteins are shown in **Fig. 1**.

**Figure 1.**
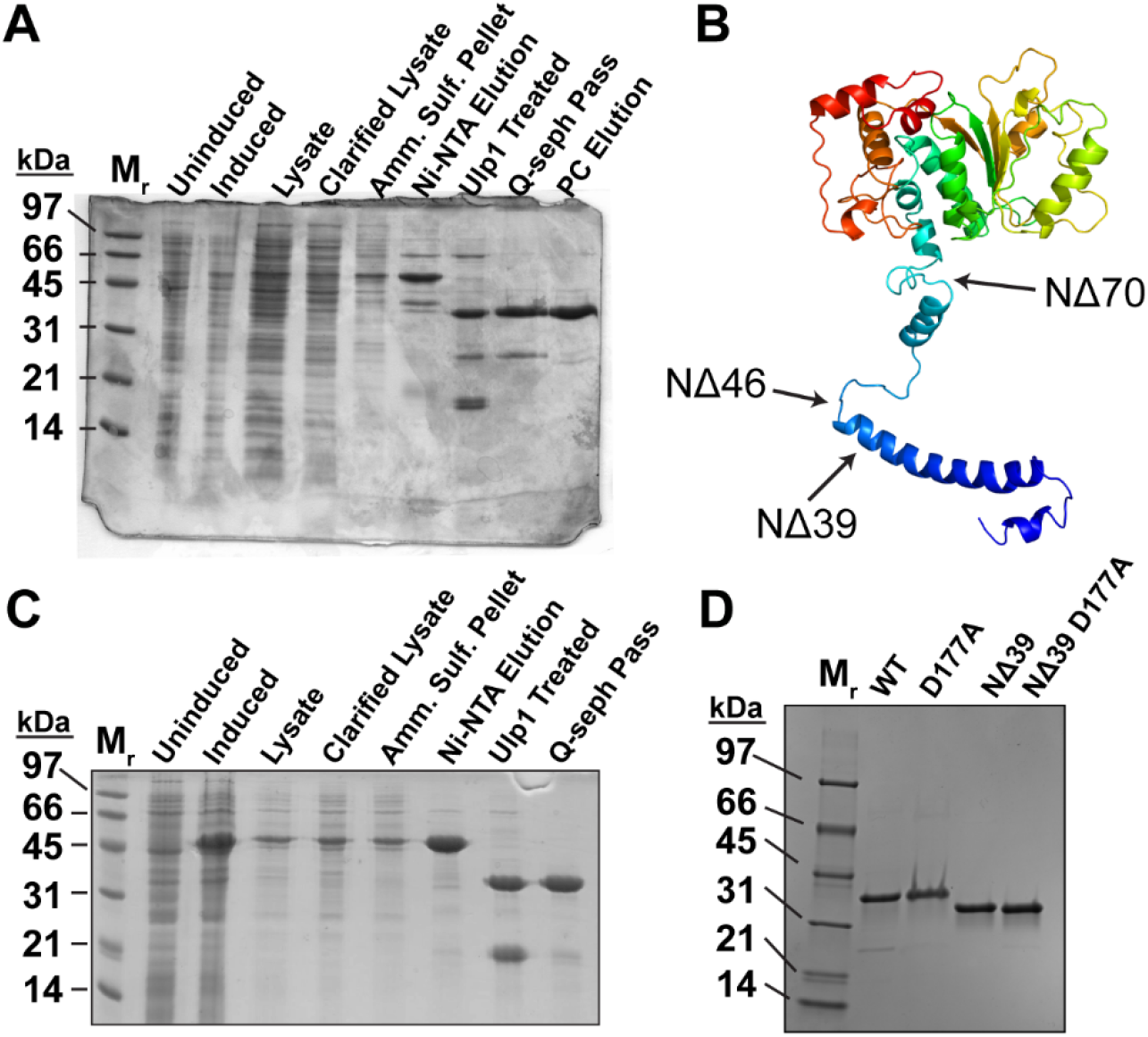
Purified PV 2C protein and derivatives used in this study. **(A)** Purification of PV 2C full-length protein (WT). We expressed and purified WT and derivatives as described under Materials and Methods. Progress was visualized before and after each step by electrophoresis through a 15% polyacrylamide gel. Briefly, cells were induced (uninduced and induced), disrupted and centrifuged (lysate vs. clarified lysate), and precipitated by ammonium sulfate (Amm. Sulf. Pellet). Next, pooled fractions of proteins eluted from the Ni-NTA column (Ni-NTA Elution), then treated with the Ulp1 protease to remove the His-SUMO tag (Ulp1-treated), Finally, protein was passed through a Q-sepharose column (Q-seph Elution), and then eluted from a phosphocellulose column (PC Elution). **(B)** PHYRE2-predicted structure of PV WT 2C modeled based on the structure of the ATP-γ-S-bound VAT complex (PDB: 5G4G). Blue is N-terminus; red is C-terminus. We deleted the indicated residues from 2C to produce NΔ39, NΔ46 and NΔ70. Only NΔ39 was soluble. **(C)** PV 2C NΔ39 was purified as described for WT in panel A. **(D)** Purified PV WT and NΔ39 (2 μg) used in this study. D177A represents the amino acid substitution used to inactivated the ATPase activity; D177 is in the conserved Walker-B motif. Molecular weight standards (M_r_) in panels A, B, and D are represented in kDa.

### Expression and purification of PV 2C NΔ115

The induction, lysis, and PEI/ammonium sulfate treatment of SUMO PV 2C 116-329 was performed as described for PV 2C WT but using buffer E [20 mM HEPES at pH 7.5, 10% (v/v) glycerol, 500 mM NaCl, 5 mM imidazole, 5 mM BME, 1 mM EDTA, 1.4 μg/mL pepstatin A, 1.0 μg/mL leupeptin]. The ammonium sulfate pellet was resuspended in Buffer F [Buffer B lacking NP-40] and containing 5 mM imidazole. Using a peristaltic pump, an Ni-NTA resin was equilibrated with 5 CVs of Buffer F containing 5 mM imidazole at 1 mL/min. The equilibrated Ni-NTA resin was then mixed with the resuspended protein pellet for 30 min while stirring, and subsequently repacked. To remove contaminants, the loaded resin was washed with 5 CVs Buffer F containing 5 mM imidazole and 4 CVs Buffer B containing 50 mM imidazole. PV 2C Δ115 was eluted into multiple fractions with Buffer F containing 500 mM imidazole. 2C protein fractions were pooled, treated with 1 μg ULP-1 per 1 mg protein of interest, and dialyzed against Buffer G [20 mM HEPES at pH 7.5, 20% (v/v) glycerol, 5 mM BME, 500 mM NaCl] overnight at 4 °C by using a 6-8,000 MWCO dialysis membrane (Spectrum Laboratories). The dialyzed 2C protein was centrifuged at 75,000 x *g* at 4 °C for 30 min to isolate soluble protein and concentrated using a Vivaspin Turbo 15 (Sartorius, 10 kDa MWCO) to 2 – 5 mL for preparative gel filtration, using a HiLoad 16/600 Superdex 200 pg (GE Healthcare). The gel filtration column was flowed at 1.0 mL min^-1^ buffer G, and divided into 34 3-mL fractions. Fractions with the protein of interest were detected by Nanodrop and SDS-Page, and subsequently pooled and concentrated with a Vivaspin Turbo 15. The protein concentration, conductivity, concentration of contaminant nucleic acids, and storage was performed similarly to PV 2C WT (PV 2C Δ115 ε_max_= 7,450 M^-1^·cm^-1^).

### Fluorescence polarization experiments

To test interactions between 2C and nucleic acids, increasing concentrations of 2C were added into a solution containing 10 nM of fluorescein-labeled nucleic acid in a binding buffer [20 mM HEPES, 5 mM magnesium acetate, 1 mM TCEP, and 25 mM NaCl, at pH 6.8] in a 100 μL final reaction volume. Reactions were incubated for 15 minutes at 30 °C and then measured. Experiments were performed in a 96-well plate format using a BioTek H1M1 plate reader using a dual wavelength 485/20 485/20 fluorescence polarization cube. Data from protein titration experiments were fit to a hyperbola:

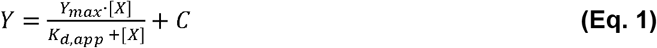

where X is the concentration of protein, Y is degree of polarization, *K*_d,app_ is the apparent dissociation constant, and Y_max_ is the maximum value of Y.

The stoichiometric binding experiment was performed essentially as described above with the addition of 3 μM unlabeled ssRNA-1. The molar ratio of 2C protein to ssRNA-1 was solved via simultaneous equations as described in the legend of **Fig. 4**.

**Figure 2.**
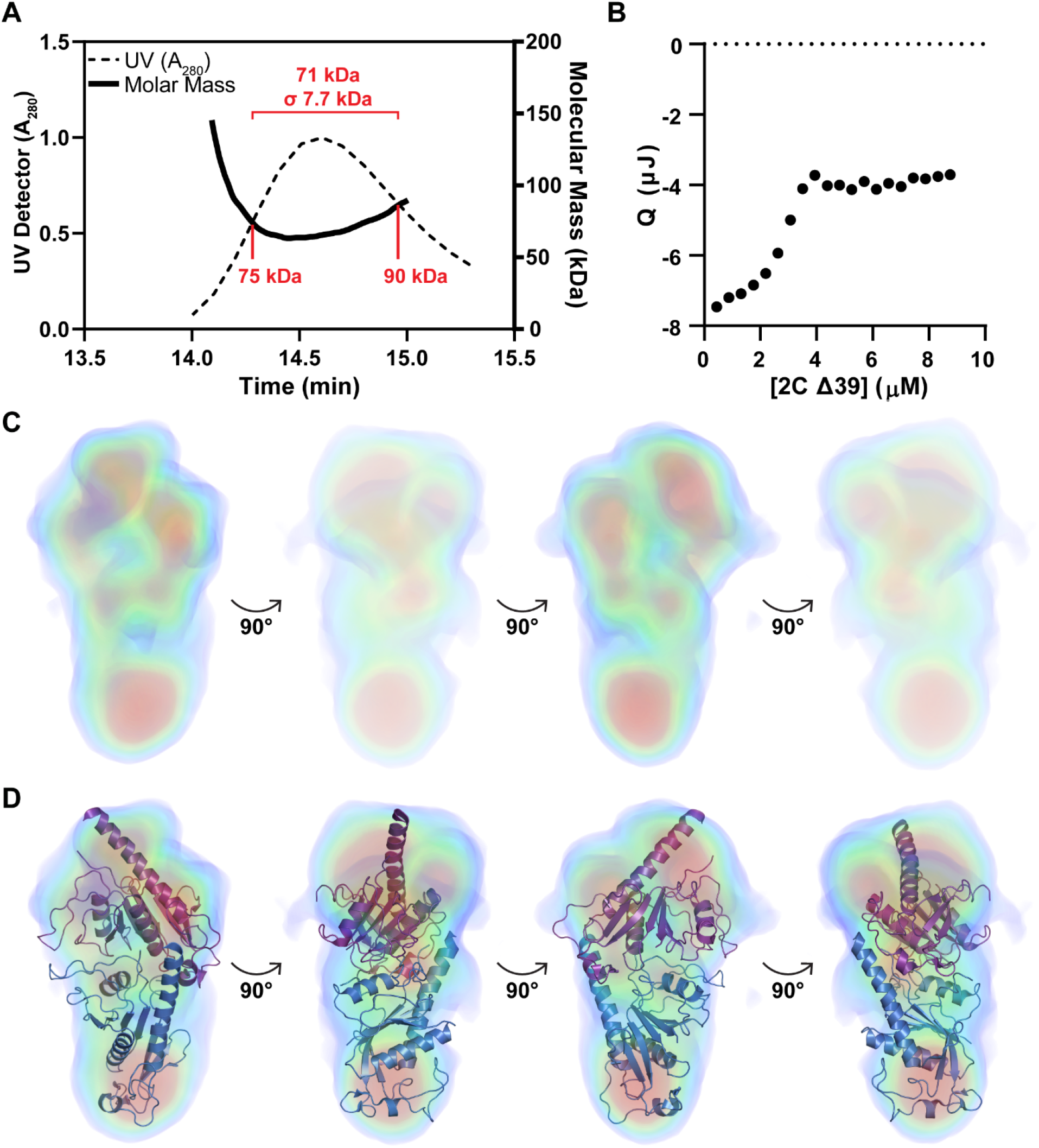
NΔ39-2C forms a dimer in solution. **(A)** Analysis NΔ39-2C by size-exclusion chromatography coupled to multi-angle light scattering (SEC-MALS) was performed as described under Materials and Methods. The sample loaded was 150 μM (5 mg/mL), with the concentration ranging from 35-50 μM during elution. Under these conditions, the average mass observed was 71 ± 7.1 kDa (95% CI: 56.8 – 85.2 kDa), most consistent with formation of a dimer (66 kDa). **(B)** Analysis NΔ39-2C by dilution isothermal titration calorimetry (ITC) was performed as described under Materials and Methods. Shown is the observed heat change as a function of the protein concentration in the cell post-injection. The data could not be fit to a model for dimer dissociation only but required addition of pre- and/or post-dissociation conformational changes. Therefore, we have not fit the data to any model. What can be gleaned from the data is that the dimer fails to dissociate at concentrations above 4 μM, suggesting a *K*_d_ value on the order of 0.4 μM. **(C)** Analysis of NΔ39-2C by small-angle X-ray scattering (SAXS) was performed as described under materials and methods. The primary data are presented in **Fig. S3**. Shown here is the density map calculated by Density from Solution Scattering (DENSS) algorithm and rotated 360 degrees (40). **(D)** Structural model of NΔ115-2C presented in **Fig. 3** fits into the density observed for NΔ39-2C by SAXS.

**Figure 3.**
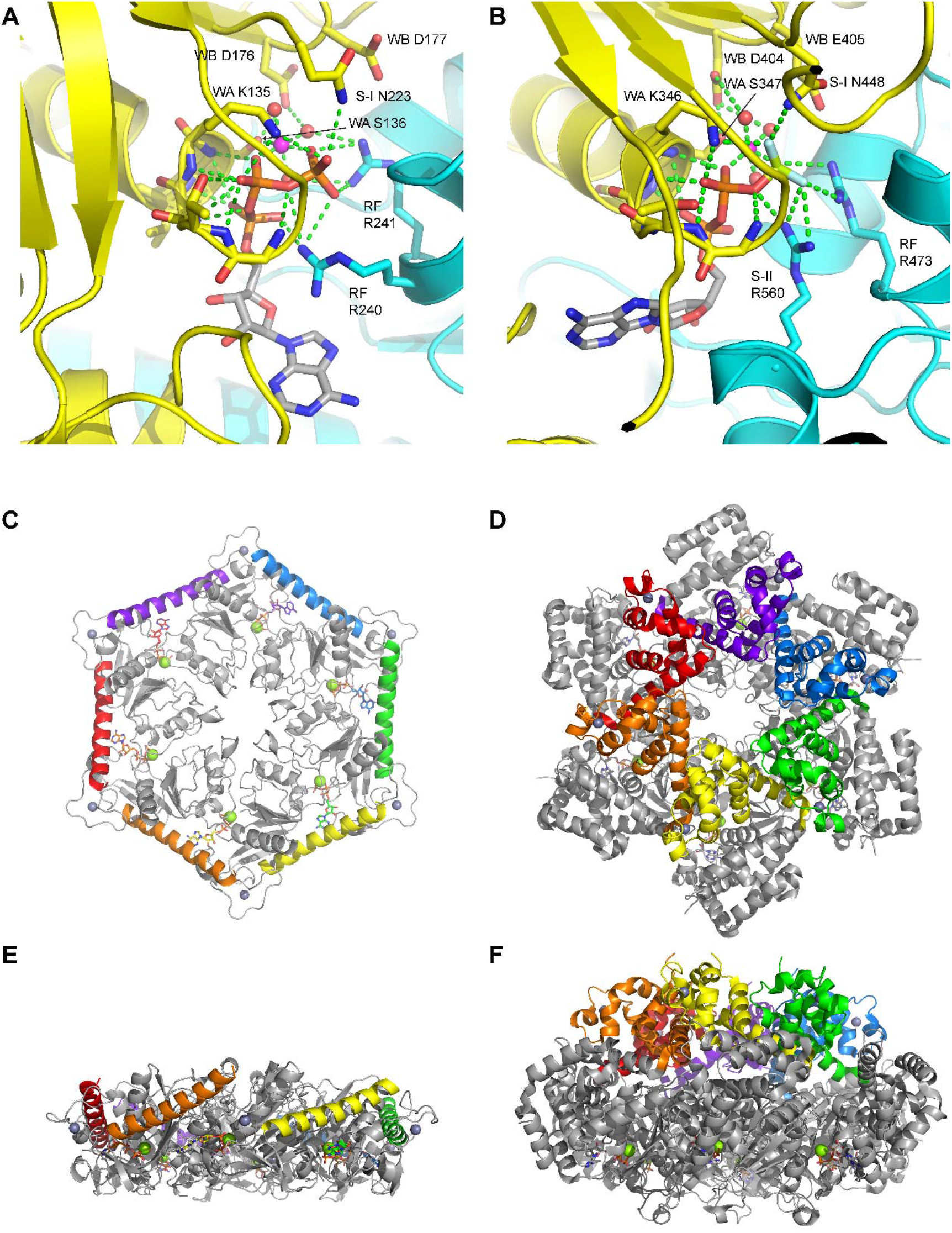
Modeling of the PV NΔ115-2C dimer. **(A)** One ATPase site from the generated hexamer model of 2C. **(B)** A similar ATPase site from MCM (PDB 6MII) **(C)** Overall 2C hexamer looking down the channel. Everything is in grey except the helix of oligomerization, which is colored by subunit. **(D)** A similar view of SV40 T-antigen. Everything is in grey except the oligomerization domain, which is colored by subunit. **(E)** A perpendicular view of panel C. **(F):** A perpendicular view of Panel D.

**Figure 4.**
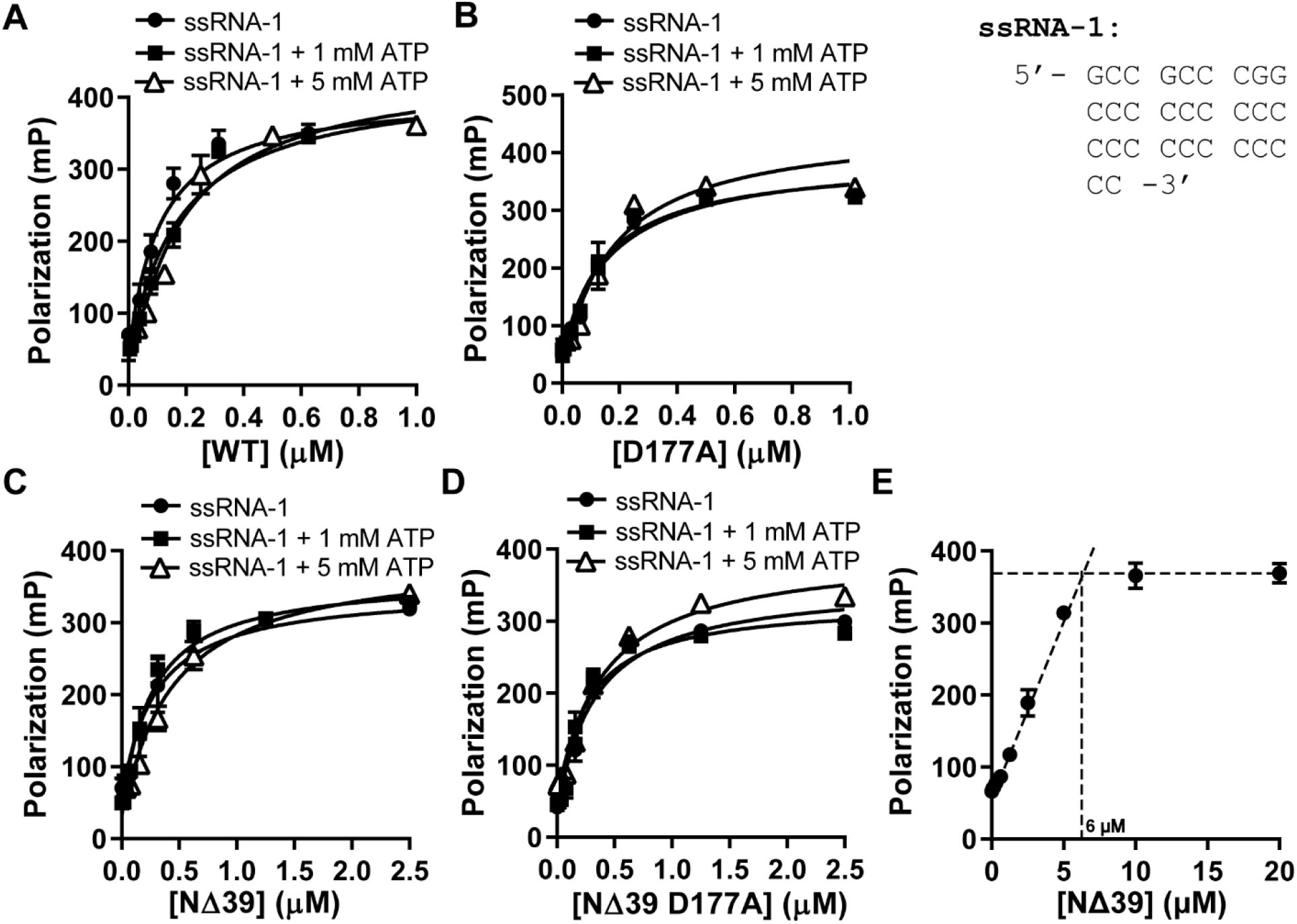
2C protein binds RNA as a dimer. **(A-E)** 2C binds to ssRNA in the absence and presence of ATP. Fluorescein-labeled ssRNA-1 (10 nM; 29-mer; see **Table S1** for additional details) was mixed with increasing concentrations of 2C (WT, NΔ39, D177A, or NΔ39 D177A) in the absence or presence of ATP (500 μM). Milliporization (mP) was plotted and the data were fit to Equation 1, yielding the apparent dissociation constants (*K*_d,app_) summarized in **Table 1**. Error bars represent the SD (n = 3). (**E**) Stoichiometry of binding between NΔ39 2C and ssRNA-1. NΔ39 2C was titrated into the RNA binding assay in which the total concentration of ssRNA-1 was 10·*K*_d,app_ (3 μM), and the mP was measured. Error bars represent the SD (n = 3). We defined lines for the two phases of the curve by linear regression and calculated the intersection (6 μM) algebraically. This value is twice the concentration of RNA indicating that 2C binds RNA as a dimer.

### Radiolabeled ATPase Assay

Reactions were performed at 30 °C and contained 20 mM HEPES pH 6.8, 25 mM NaCl, 5 mM magnesium acetate, 0.05 μM [α^32^P]-ATP, and 1 mM TCEP (17). Depending on the assay, the presence of ATP, RNA, or drug varied, and are indicated within figures and their legends. Nucleic acids used for stimulation experiments are listed in **Table S1**. Reactions were quenched by the addition of EDTA at a final concentration of 250 mM. During nucleotide competition experiments, the total amount of Mg^2+^ was stoichiometric compared to the total NTP concentration, with an additional 4 mM Mg^2+^ free. The volume of enzyme added to any reaction was always less than or equal to one–tenth the total reaction volume. Rates are defined as μM ADP formed min^-1^ μM 2C^-1^. When defining the rate, timepoints were taken during the linear portion of the reaction with less than 20% product formed.

### Radiolabeled ATPase assay product analysis

Products were resolved by spotting 1 μL of quenched reactions onto polyethyleneimine (PEI) TLC plates (Fisher Scientific M1055790001). TLC plates were developed in 0.3 M potassium phosphate, pH 7.0, dried, and exposed to a PhosphorImager screen. Phosphor images were taken with a GE Amersham Typhoon. Products were quantified using ImageQuant software (Molecular Dynamics) to determine the amount of ADP formed from ATP. For linear reactions, the amount of ADP formed was plotted as a function of time and fit to a line. For saturating reactions, the rate (defined as μM ADP min^-1^ μM 2C^-1^) was plotted versus concentration of substrate and fit to a hyperbola:

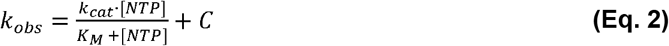

where [NTP] is the concentration of ligand, *k_obs_* is the rate, *k_cat_* is the maximal rate, K_M_ is the Michaelis constant, and C is a constant. When fitting data from competition experiments, the following equation was used:

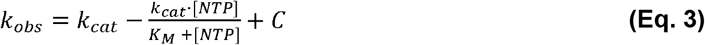

When necessary, a quadratic binding equation was used to provide another perspective where the free concentration of protein was not assumed to be the concentration of dimer in solution. To calculate the concentration of monomer, a quadratic was used:

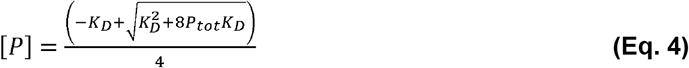

where [P] is the concentration of monomer, P_tot_ is the total protein concentration, and *K_D_* is the equilibrium dissociation constant of the dimer. Eq. 4 was then used to calculate the concentration of dimer:

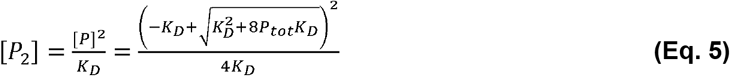

where [P_2_] is the concentration of dimer. Finally, concentration dependent rate (defined as μM ADP min^-1^ μM 2C^-1^) activity curves were fit with:

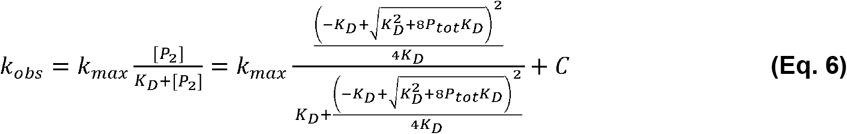

where *k*_max_ is the maximum rate with a substrate concentration of 0.5 mM ATP, C is a constant, and *k*_obs_ the observed rate (μM ADP min^-1^ μM 2C^-1^).

### Colorimetric ATPase assay

The BioAssay Systems QuantiChrom ATPase/GTPase Assay Kit (BioAssay Systems, Hayward, CA, USA) was used for colorimetric ATPase assays. Reactions were performed at 30 °C and contained 20 mM HEPES pH 6.8, 25 mM NaCl, 5 mM magnesium acetate, NTP, and 1 mM TCEP. Depending on the assay, the presence of NTP, or RNA varied, and are indicated within figures and their legends. Timepoints were taken by quenching with 50 mM final EDTA. 20 μL of quenched reaction mix was added to 80 μL of QuantiChrom reagent in a 384-well glass-bottom plate and left to develop for 30 minutes. The A_620_ was then read using a BioTek H1M1 plate reader. A phosphate standard curve was produced using the provided phosphate standard in the same reaction buffer used for 2C enzymatic assays and was linear between 1 and 50 μM phosphate: samples were diluted into that range for final measurement. The standard curve was used to calculate concentration of phosphate present in assay timepoints or samples.

### Deprotection and 5’-^32^P-labeling of RNA oligonucleotides

RNA nucleotides were ordered to be PAGE-purified in their ACE-protected forms and were deprotected according to instructions of the manufacturer. For helicase assays, oligonucleotides were labeled using [γ-^32^P]-ATP and T4 polynucleotide kinase as previously described (37).

### Annealing of unwinding substrates

RNA duplexes (pairs made up of ssRNA 2 and 3, ssRNA 4 and 5, and ssRNA 7 and 8; sequences detailed in **Table S1**) were annealed at 5 μM in 20 mM HEPES, pH 7.5, 50 mM NaCl by using a Progene Thermocycler (Techne). The unlabeled strand was present in a 1.2:1 ratio excess. Annealing reactions were heated to 90 °C for one minute and cooled 5 °C min^-1^ until 10 °C.

### Helicase assay

Reactions (20 μL) were performed at 30 °C with either RNA duplex. We used either Mix A or Mix B. Mix A contained 20 mM HEPES pH 6.8, 25 mM NaCl, 5 mM magnesium acetate, 1 mM ATP, 5 μM DNA trap strand, and 1 mM TCEP. Mix B contained 20 mM HEPES pH 7.0, 25 mM NaCl, 0.5 mM MgCl_2_, 2 mM ATP, 5 μM DNA trap strand, and 1 mM TCEP (38). The trap strand was a 9-mer RNA of the same sequence as the ^32^P-labeled strand, 5’-CCGGGCGGC-3’. Reaction timepoints were quenched with an equal volume (5 μL) addition of loading buffer (100 mM ETA, 0.33% (wt/v) SDS, 10% (v/v) glycerol, 0.025% (wt/v) bromophenol blue, 0.025% (wt/v) xylene cyanol). Products were resolved on 20% native PAGE gels by electrophoresis in 1x TBE at 15 mA for 90 minutes. Gels were visualized by phosphorimaging.

### Western Blotting of 2C protein

Rabbit polyclonal antibody against 2C was produced by Covance Research Products (Denver, PA) using the antigen PV 2C NΔ115. 100 picograms (100 pg) of purified 2C WT and NΔ39 protein was loaded onto a 15% SDS PAGE gel. Mock, 4, and 6 hour post-infection (hpi) HeLa cell extracts (60 x 10^3^ cells lane^-1^) were loaded onto the same gel. After electrophoresis, the SDS-PAGE gel was sandwiched to a nitrocellulose membrane with a Genie Blot apparatus (IDEA Scientific Company). The membrane was probed with 1:5000-dilution of the final bleed of anti-2C polyclonal antibody, and 1:5000-dilution of goat anti-rabbit IgG-HRP (Santa Cruz Biotechnology). The 2C-reactive bands were detected by ECL Western blot detection reagents (Azure Biosystems) and analyzed by a G:Box imager (SynGene).

### Size-exclusion Chromatography and Multi-angle Light Scattering (SEC-MALS)

Samples of 2C NΔ37 protein at a concentration of 5mg/ml were prepared in a buffer of 20 mM HEPES, pH 6.8, 5 mM Magnesium acetate, 1 mM TCEP, 5% glycerol, and 50 mM NaCl. All samples were centrifuged at 6,000 x *g* for 20 min to minimize aggregation and to remove dust particles prior to SEC-MALS. SEC chromatographic separation of samples was conducted using an Agilent 1260 Infinity II HPLC system with an autosampler and fraction collector. A Wyatt Technology DAWN MALS and Wyatt Optilab Refractive Index(RI) detectors were used for analyzing the molar mass of peaks that eluted from the column. The SEC-MALS system was calibrated with bovine serum albumin (BSA) and equilibrated with in the same mobile phase as that of the 2C samples. Normalization and alignment of the MALS and RI detectors were carried out using the BSA (monomer ~66 kDa) standard, run in the 2C buffer condition. The Wyatt SEC hydrophilic column used had 5um silica beads, a pore size of 100 Angstrom and dimensions 7.8 x 300 mm. The column, fraction collector and autosampler were set to 4°C. A volume of 15 μL of 2C protein (75 μg) was injected at a flow rate of 0.5 mL min^-1^ with a chromatogram run time of ~25 minutes. Data was analyzed using the ASTRA software from Wyatt. As a single peak with a molar mass corresponding to a dimer of 2C protein was observed by SEC-MALS, BioSAXS data were subsequently collected directly with no further purification via SEC-MALS fractions.

### BioSAXS and modeling of the protein envelope

Small angle X-ray scattering (BioSAXS) was collected on 2C NΔ39 (at 2 mg ml^-1^ and 1 mg ml^-1^) and 2C NΔ39 with ATP (at 1 mg ml^-1^) to compare their solution state conformations. BioSAXS data were collected at a wavelength of 1.54 Å on the in-house home source X-rays generated by a Rigaku MM007 rotating anode housed with the BioSAXS2000nano Kratky camera system. The system includes OptiSAXS confocal max-flux optics that is designed specifically for SAXS and a sensitive HyPix-3000 Hybrid Photon Counting detector. The sample capillary-to-detector distance was 495.5 mm and was calibrated using silver behenate powder (The Gem Dugout, State College, PA). The useful q-space range (4πSinθ λ^-1^, *2Θ*being the scattering angle) was generally from qmin= 0.008 Å^-1^ to qmax= 0.3Å^-1^ (q = 4ττsin(θ) λ^-1^, where *2Θ* is the scattering angle). The energy of the X-ray beam was 1.2 keV, with the Kratky block attenuation of 22% and a beam diameter of ~100 μm. Protein samples were loaded using the autosampler on to quartz capillary flow cell mounted on a stage cooled to 4 °C prior and aligned in the X-ray beam. The sample cell and full X-ray flight path, including beam stop, were kept in vacuo (< 1×10^-3^ torr) to eliminate air scatter. The Rigaku SAXSLAB software was programmed for automated data collection of each protein with elaborate cleaning between samples. Data reduction including image integration and normalization, and background buffer data subtraction were also carried out using the SAXSLAB software. Six ten-minute images from protein and buffer samples were collected and averaged after ensuring that no X-ray radiation damage had occurred. SAXS data overlays showed that there was no radiation decay over the 60 minutes of data collection. Data analysis was done in ATSAS and solvent envelopes were computed using DENSS, an algorithm used for calculating ab initio electron density maps directly from solution scattering data (39,40). The theoretical scattering profiles of the constructed models were calculated and fitted to experimental scattering data using CRYSOL.

### Modeling of the PV NΔ115-2C dimer

To construct a model for the PV NΔ115-2C dimer, we sought to recapitulate likely features of the bipartite AAA+ ATPase site and maintain compatibility with a hexameric arrangement, which is adopted in the helicase-active forms of other SF3 helicases such as papillomavirus E1 (41,42) and SV40 Large T-antigen (43,44). Similar to the procedure applied previously for modeling of the PV 2C hexamer (45), a hexameric model for a NΔ115-2C hexamer was constructed by superposition of the structure of the 2C ATPase domain (chain A, PDB 5Z3Q (45)) onto the AAA+ domains of an SV40 Large T-antigen hexamer. To facilitate further inspection of the ATPase sites, the structure of SV40 Large T-antigen hexamer bound to 6 ATP molecules and 6 magnesium ions was used (chains A-F, PDB 1SVM) rather than the structure used previously (45) of T-antigen bound to p53 (PDB 2H1L (46)), which lacks magnesium and nucleotide. Six copies of the 2C AAA+ domain were superimposed on those of SV40 Large T-antigen in Coot (47) using the secondary structure matching algorithm.

Following this superposition, the positions of the ATP, magnesium, and three water molecules bound to the magnesium from chain A of the SV40 Large T-antigen structure (1svm chain A, waters 802, 935, 936 (43)) were optimized for alignment to the Walker-A motif of one 2C subunit by explicit superimposition of the Walker-A helix of both structures (aligning residues 429-439 of 1SVM to residues 132-142 of the superimposed 2C subunit; see above). Following this adjustment, the Mg·ATP·water unit was examined and revealed a substantial steric clash of the adenine ring with the 2C protein. The adenine ring was rotated to an alternate non-clashing conformation in Coot (47) and minimized. This minimized Mg·ATP·water unit was copied to the other subunits of the putative 2C hexamer model by least-squares alignment of the ATP-containing subunit to each of the remaining five subunits to generate a hexamer with ATP and an octahedrally coordinated magnesium at all six ATPase sites. Finally, the overall hexameric structure was minimized in Phenix (48) with phenix.geometry_minimization.

This minimization incorporated distance restraints for likely interactions of the ATPase site, guided by the structure of the MCM AAA+ hexameric helicase (PDB 6MII (49)). The specific restraints included an octahedral coordination sphere (1.95 Å bond distances and 90° bond angles) for the magnesium ion comprised by the conserved hydroxyl of the Walker-A serine (S136), an ATP β-phosphate oxygen, an ATP γ-phosphate oxygen, and three water molecules. Some hydrogen-bonding distance restraints (3.0 Å) were applied between protein atoms and the ATP molecule, detailed further below. The main-chain amide groups of Walker-A residues G132-V137 and oxygen atoms of ATP phosphates were restrained to 3.0 Å. The side-chain ammonium of the conserved lysine of the Walker-A motif (K135) and oxygen atoms of the ATP were restrained to 3.0 Å. The sensor-I conserved asparagine (N223) side-chain and an ATP oxygen atom were restrained to 3.0 Å. From the neighboring subunit, two arginine residues, the arginine finger (R241) and its neighbor (R240) and oxygen atoms of the ATP phosphates were restrained to 3.0 Å. Based on comparison with the MCM ATPase site, the R240 side-chain appears to fulfill a role equivalent to sensor-II (2C does not have a canonical sensor-II sequence motif, consistent with other SF3 helicases such as E1, which places a lysine side-chain in the structural position typically occupied by a sensor-II arginine (41)). A conserved aspartate of the Walker-B motif (D176) was restrained to 3.0 Å from a water molecule of the magnesium coordination sphere and the conserved hydroxyl of the Walker-A motif (S136). The resulting hexamer model showed a good hexameric geometry free of clashes and six equivalent ATPase sites with interactions between the protein, ATP, and magnesium that are anticipated to be important for catalysis. Two adjacent subunits from this model were selected as the PV NΔ115-2C dimer model.

### Isothermal Titration Calorimetry (ITC)

Dilution experiments were performed using a TA instruments Affinity ITC auto equipment (TA instruments New Castle, Delaware). Protein PV 2C (27 μM) was loaded onto the titration syringe. The calorimeter cell contained exactly matched buffer: 10 mM HEPES, 50 mM NaCl, 5% glycerol, 1 mM TCEP, at pH 7.5. Enthalpy of dimer dissociation were recorded with injection volume of 3 μL over 20 titrations. All the injections were conducted at a constant temperature of 30 °C with a stir speed of 125 rpm. The ITC dilution data were integrated and analyzed using the TA Instruments Nano analyze software program (v3.12.0). Before testing protein samples, the instrument was calibrated using the EDTA/CaCl_2_ test kit for the TA Affinity ITC auto.

### Quantification and Statistical Analysis

Statistical analysis and nonlinear regression was performed using GraphPad Prism v9.3. Error bars in graphs represent standard deviation. Plus-minus values provided in tables and legends are the standard error. An unpaired Student’s t-test was used to assess stimulation of 2C by double-stranded RNA. The P values for the experiment are presented in the legend (n.s. = non-significant, * = < 0.05, ** = <0.005, *** = < 0.005).

## RESULTS

### Expression and purification of PV 2C protein and derivatives

The enteroviral 2C protein has been notoriously difficult to purify (17,27–29,50). The presence of an amino-terminal, amphipathic alpha-helix on the protein causes aggregation and formation of inclusion bodies when expressed in E. coli (17,45,51). Previous studies of the NS5A protein from hepatitis C virus showed that fusion to cleavable tags positively augments expression and solubility (45,51). We have used a similar strategy for expression of PV 2C protein. We encoded a six-histidine-SUMO-tag on the amino terminus (30). We induced the protein using the auto-induction system, which leads to expression of chaperones capable of preventing protein aggregation (35). The combination of these two approaches permitted purification of the authentic, full-length 2C protein as described under Materials and Methods (**Fig. 1A**). We will refer to a protein with an authentic amino terminus as WT. Solubility of WT, full-length 2C requires the presence of detergent.

Structural information is available for a derivative of PV 2C in which the amino-terminal 115 amino acid residues were deleted (45,51). We will refer to this protein as NΔ115. We purified NΔ115; however, it lacked ATPase activity under conditions used for WT 2C (**Fig. S1**). Modeling of PV 2C based on a structure for the VAT AAA+ ATPase placed amino acid 115 well into the body of the folded globular domain of 2C (**Fig. 1B**). Therefore, we engineered several shorter truncations with the goal of identifying a derivative that was soluble in the absence of detergent and active. Deleting 79 or 46 amino acid residues from the amino terminus produced insoluble proteins (data not shown). However, deletion of only 39 amino acid residues produced a soluble protein that we purified using the protocol developed for WT, but without the requirement of detergent for solubility (**Fig. 1C**).

The Xia laboratory has reported the existence of post-translational modification on picornaviral 2C proteins expressed in insect cells that alter the mobility of the protein on denaturing gels (26). This modification is also thought to be required for helicase activity (26). Interestingly, WT PV 2C proteins produced in bacteria or during infection exhibit the same mobility on denaturing gels (**Fig. S1D**). Recently, the Cui laboratory expressed a picornaviral 2C protein in insect cells (52). In this case, perturbed gel mobility was not observed (52).

To inactivate the ATPase activity of 2C without impairing nucleotide binding, we changed the conserved Asp-177 to Ala (D177A). Asp-177 is a residue in the “Walker B motif,” which contributes to formation of the nucleophile in the active site (**Fig. S2A**) (53,54). Purification of the D177A variants in the context of the full-length or NΔ39 proteins followed protocols developed for the unsubstituted enzymes. As expected, the D177A substitution inactivated the ATPase activity (**Fig. S2B,C**). Purified proteins used in this study are shown in **Fig. 1D**.

### PV 2C-NΔ39 exists as a dimer in solution

SF3 helicases are active as hexamers. Previous studies of picornaviral 2C proteins have demonstrated the capacity of these proteins to form hexamers. However, the efficiency with which hexamers form is very low (20,28,29). Our ability to purify the NΔ39 derivative in the absence of detergent and to concentrate the protein to a concentration of 5 mg/mL (150 μM) or greater provided the opportunity to use a variety of biophysical approaches to characterize the behavior of NΔ39 in solution properties.

We evaluated the quaternary structure of NΔ39 using size-exclusion chromatography coupled to multiangle light scattering (SEC-MALS) as described under Materials and Methods. Neither monomer nor hexamer formed in solution based on this experiment (**Fig. 2A**). The average mass of the species present in solution was 71.0 ± 7.7 kDa (**Fig. 2A**). Given the calculated molecular weight of 33 kDa for a monomer, these data are most consistent with NΔ39 existing as a dimer in solution. The concentration range of protein eluted from the column during the SEC-MALS experiment was 35 – 50 μM based on the UV absorbance profile (**Fig. 2A**). Use of high protein concentrations is a requirement for this experiment, so it is not possible to glean any insight into the equilibrium dissociation constant for the dimer from this experiment. However, the inability to observe hexamers over this range of concentration suggests that the equilibrium dissociation constant for such structures must be quite high.

To pursue information on the equilibrium dissociation constant (*K*_d_) for the dimer, we used isothermal titration calorimetry (ITC). We used a dilution format as described under Materials and Methods. The basis for this experiment is that if the protein exists as a dimer in solution, then dilution of the protein into buffer to a concentration below the value of the *K*_d_ will cause a heat change due to dissociation of the dimer. Such an effect will be observed until the concentration of protein in the cell exceeds the *K*_d_ value. This phenomenon was observed for NΔ39, consistent with the existence of a dimer in solution (**Fig. 2B**). Unfortunately, we were unable to model the data to a simple dimer-to-monomer transition to obtain a value for the *K*_d_. However, it is clear that concentrations above 4 μM failed to exhibit heat change (**Fig. 2B**). If we assume that heat changes will be undetectable when the protein concentration in the cell reaches a value of 10·*K*_d_, then the *K*_d_ value for the dimer would be on the order of 400 nM.

Finally, to evaluate the quaternary structure of 2C further and to reveal the overall three-dimensional shape of the protein in solution, we performed small-angle X-ray scattering (SAXS). The threedimensional protein envelope calculated using the SAXS data is shown in **Fig. 2C**. Kratky plots confirmed a well folded protein with no flexibility or disorder (**Fig. S3**). Values for the radius of gyration (24.1 Å) and maximum radius (D_max_ of 86 Å), as well as values from the distance distribution (P(r)), CRYSOL, and Guinear analyses, were all consistent with the 2C dimer as the predominant species in solution.

### Structural model for the PV 2C hexamer based on the NΔ115-2C structure

We used the structure of PV NΔ115-2C to construct a model of the PV 2C hexamer as described under Materials and Methods. Modeling was templated by SV40 Large T-antigen, an SF3 helicase, bound to ATP and Mg^2+^ (PDB 1SVM) (44). Modeling of the organization of the active site at each dimer interface of the NΔ115-2C hexamer was restrained by the structure of the MCM AAA+ hexameric helicase solved in the presence of ATP and Mg^2+^ (PDB 6MII) (49). The final model of the NΔ115-2C hexamer included ATP and Mg^2+^ bound to each dimer interface of the hexamer. A single dimer interface is shown in **Fig. 3A**. The Mg^2+^ ion exhibited an octahedral coordination sphere. Ligands to the metal included the following: the conserved hydroxyl of Walker-A serine (S136), oxygens from the β- and γ-phosphates of ATP, as well as several water molecules (**Fig. 3A**). The phosphates of ATP interacted with the conserved Walker-A lysine (K135), the conserved sensor-I asparagine (N223), the conserved arginine-finger arginine (R241), and R240 (**Fig. 3A**). One of the conserved Walker-B aspartates (D176) interacted with both the Mg^2+^ ion and the hydroxyl of the conserved Walker-A serine (**Fig. 3A**). The other aspartate (D177) had no direct contact with the nucleotide (**Fig. 3A**), yet is essential for catalysis (51,55,56). The organization of the modeled 2C active site was consistent with the active site observed in the structure of MCM in complex with ATP and Mg^2+^ (**Fig. 3B**) (49). Interestingly, R240 of 2C may serve the same function as the conserved sensor-II arginine (R560) of MCM (compare **Fig. 3A** to **Fig. 3B**). Importantly, the dimer modeled here fit nicely into the solution envelope determined experimentally SAXS above (**Fig. 2D**).

As reported by the Cui laboratory, enteroviral 2C monomers interact by insertion of the carboxy-terminal helix, referred to as the pocket-binding motif, of one monomer into hydrophobic pocket of another (45,51). We modeled our PV 2C hexamer to contain the same interactions (**Fig. 3C**). Such head-to-tail interactions could extend ad infinitum. The Cui laboratory has shown that this is not the case in solution, as species no larger than tetramer are observed in solution for PV NΔ115-2C (45,51). As reported above, we did not observe species larger than dimer in solution. How 2C-2C quaternary structure is regulated remains unclear.

We compared the PV 2C hexamer to that of SV40 Large T-antigen. For T-antigen, a domain comprised of multiple helices contributes to stabilization of the hexamer (**Fig. 3D**). By comparing side views of the 2C hexamer (**Fig. 3E**) to that of T-antigen (**Fig. 3F**), it becomes even clearer that T-antigen uses an entire domain to hold the subunits of the hexamer together. The solvent accessible surface area buried in forming the T-antigen hexamer is on the order of 2600 Å^2^, but only 1200 Å^2^ for the 2C hexamer. Determinants other than the interactions driven by the carboxy-terminal helix of 2C may be required to form a stable hexamer.

### A PV 2C dimer binds to RNA

Although there has been some suggestion that PV 2C binds viral RNA (57,58), quantitative information on the affinity and stoichiometry of binding does not exist. We have used a fluorescence-polarization assay to monitor RNA binding as described under Materials and Methods. We titrated WT protein into binding reactions in the absence or presence of ATP. We plotted polarization as a function of WT protein concentration and fit the data to a cooperative-binding mechanism (see Materials and Methods), yielding an apparent dissociation constant (*K*_d,app_). WT bound to RNA with a *K*_d,app_ of 110 ± 20 nM (**Fig. 4A**). The presence of ATP caused an insignificant decrease in binding affinity (**Fig. 4A**). NΔ39 was also capable of binding RNA with a *K*_d,app_ of 320 ± 60 nM (**Fig. 4B**), without a significant change caused by the presence of ATP. The values measured for NΔ39 were consistent for multiple preparations of this derivative (**Fig. S4**). To better assess the range of data points used for quantitative analysis, data have also been plotted on a log scale (**Fig. S5**).

The motivation for evaluating the impact of ATP on RNA binding comes from our previous studies of the HCV ATPase/RNA helicase that suggest ATP binding loosens the grip of the enzyme on RNA. Because WT will hydrolyze ATP, it was possible that the magnitude of the effect of ATP on RNA binding by 2C was reduced. To address this possibility, we evaluated the D177A enzyme. In each case, the inactive 2C protein bound to RNA with a similar affinity as WT or NΔ39 but without any discernible impact of ATP on binding to RNA (**Figs. 4C,D** and **S5**). These data are summarized in **Table 1.**

**Table 1.** Summary of nucleic-acid binding to 2C in the presence and absence of ATP.

| [ATP] | Dissociation Constant<sup>a</sup> of<br>WT 2C<sup>b</sup>, $K_{d,app}$ (nM) | | Dissociation Constant<sup>a</sup> of<br>NΔ39 2C<sup>b</sup>, $K_{d,app}$ (nM) | |
| --- | --- | --- | --- | --- |
|  | WT | D177A | NΔ39 | D177A |
| **0 mM** | 110 ± 20<br>[75 – 150] | 170 ± 30<br>[110 – 190] | 320 ± 60<br>[210 – 440] | 330 ± 50<br>[230 – 420] |
| **1 mM** | 180 ± 30<br>[125 – 230] | 150 ± 20<br>[110 – 190] | 250 ± 30<br>[180 – 310] | 240 ± 40<br>[160 – 320] |
| **5 mM** | 170 ± 30<br>[65 – 270] | 240 ± 70<br>[80 – 390] | 430 ± 70<br>[290 – 570] | 400 ± 50<br>[290 – 510] |

<sup>a</sup>The dissociation constant is presented ± standard error with 95% confidence interval provided in brackets below.
<sup>b</sup>Data from **Fig. 4**.

As indicated above, the prevailing view of the field is that picornaviral 2C proteins exist and function as hexamers (20,28,29,45,51,59), although our preparations only form dimers (**Fig. 2**). To determine the stoichiometry of binding to RNA, we elevated the concentration of RNA used in the experiment to 3 μM, which is more than 10-times the value for *K*_d,app_. For this experiment, we used NΔ39 because of its solubility in the absence of detergent. This experiment yielded a stoichiometry of two molecules of 2C, a dimer, per molecule of RNA (**Fig. 4E**).

The value for polarization measured correlates with the mass of the complex in solution. The observation of essentially the same endpoint in all three experiments is consistent with a dimer forming in all three experiments (**Figs. 4A-C**). The inability of protein concentrations as high as 20 μM to drive the equilibrium to complexes of higher mass suggests that a dimer may be the terminal state of PV 2C formed in solution (**Fig. 4C**), consistent with the biophysical experiments reported above (**Fig. 2**).

What these experiments do not reveal is whether the dissociation constant measured reflects a value for the 2C dimer, RNA-bound 2C dimer, or some combination thereof. Below, we perform experiments to distinguish between these possibilities.

### PV 2C binding to nucleic acid is driven by the phosphodiester backbone

Members of SF3 include both RNA and DNA helicases. Whether specificity exists for one type of nucleic acid is not clear. In addition, the extent to which substituents other than the phosphodiester backbone contribute to affinity and/or specificity of nucleic acid binding is not known. Therefore, we evaluated the nucleic acid specificity for PV WT and NΔ39 2C proteins. In addition to RNA, we employed DNA, RNA modified at the 2’-hydroxyl, and morpholino nucleic acid (MNA) (**Fig. 5A**). MNA contains a backbone that lacks the negative charge associated with the phosphate backbone of natural nucleic acid (60).

**Figure 5.**
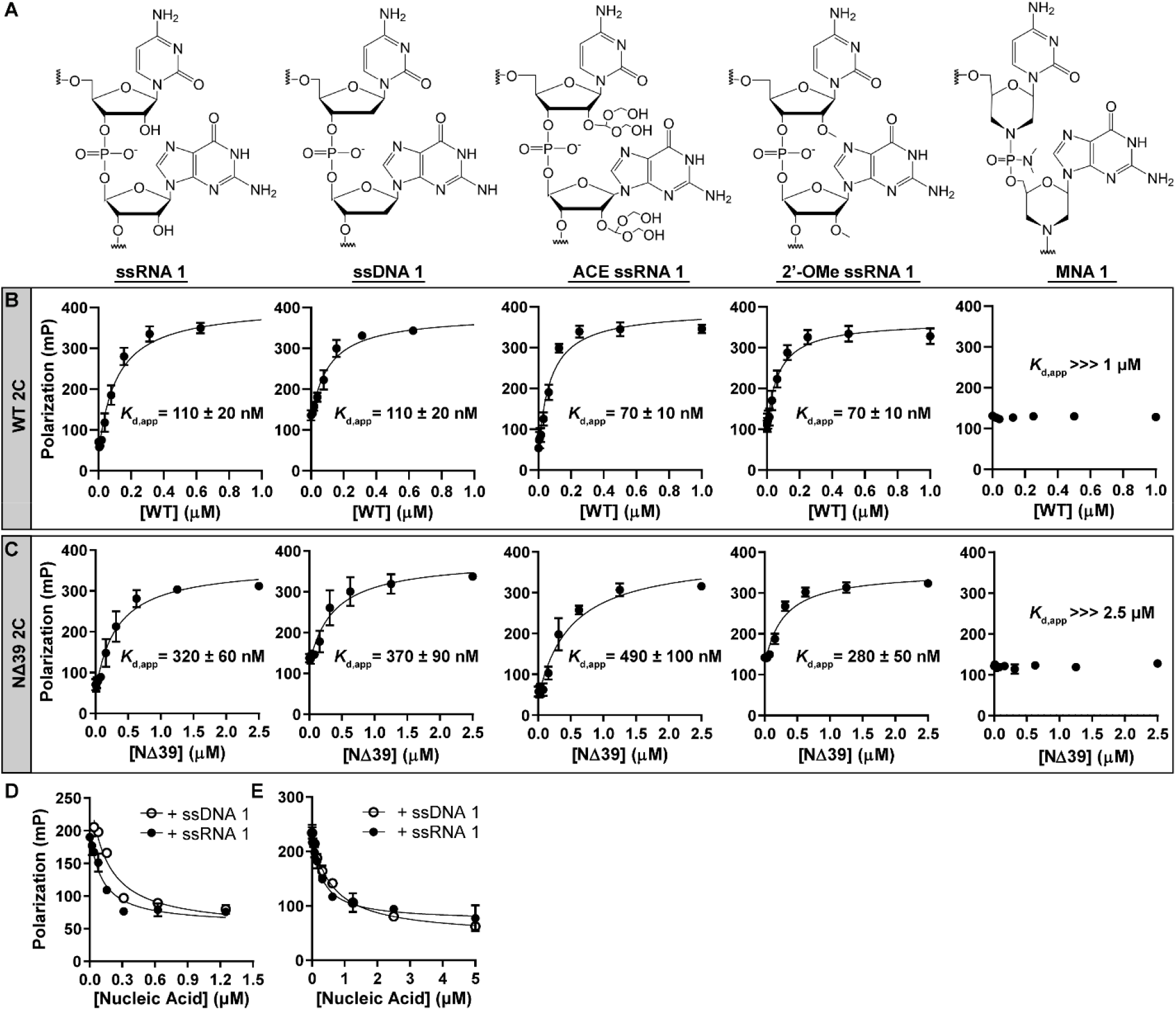
2C binding to nucleic acid is driven by the phosphodiester backbone. **(A)** Chemical structures of nucleic acids used to assess binding to 2C are as follows: ssRNA, ssDNA, 2’-ACE-protected ssRNA, 2’O-methylated (2’-OMe) ssRNA, and morpholino nucleic acid (MNA). **(B,C)** 2C-nucleic acid binding. Fluorescein-labeled nucleic acid (10 nM) was mixed with increasing concentrations of 2C (WT, panel B; NΔ39, panel C). The resulting mP was plotted and the data were fit to Equation 1, yielding the apparent dissociation constants (*K*_d,app_) and Hill coefficients (n_H_) reported in each panel. See **Table 2** for a summary of calculated values. Error bars represent the SD (n = 3). **(D,E)** 2C-RNA binding competition experiments. A fixed concentration of 2C (2 μM; WT, panel D; NΔ39, panel E) was incubated with fluorescein-labeled ssRNA-1 (10 nM) in the absence or presence of varying concentrations of unlabeled ssRNA-1 or ssDNA-1. The mP was measured and plotted. The data were fit to Equation 3. Error bars represent the SD (n = 3). We observed no significant difference in the values obtained using ssRNA-1 or ssDNA-1.

As indicated by data above (**Fig. 4C**), the affinity of NΔ39 for RNA was reduced by 3-fold relative to WT (compare RNA in **Fig. 5B** to RNA in **Fig. 5C**). Both proteins bound to DNA with affinity comparable to that of RNA (see panels DNA in **Figs. 5B,C**). Consistent with this observation, blocking the 2’-hydroxyl of RNA by addition of a 2’-acetoxy orthoester (ACE) (61) or methyl group had essentially no impact on binding (see panels ACE and 2’-OMe in **Figs. 5B,C**). In contrast, incorporation of a morpholino backbone ablated all detectable binding (see panels MNA in **Figs. 5B,C**). To better assess the range of data points used for quantitative analysis, data have also been plotted on a log scale (**Fig. S6**). Values for the dissociation constants obtained are summarized in **Table 2**.

**Table 2.**
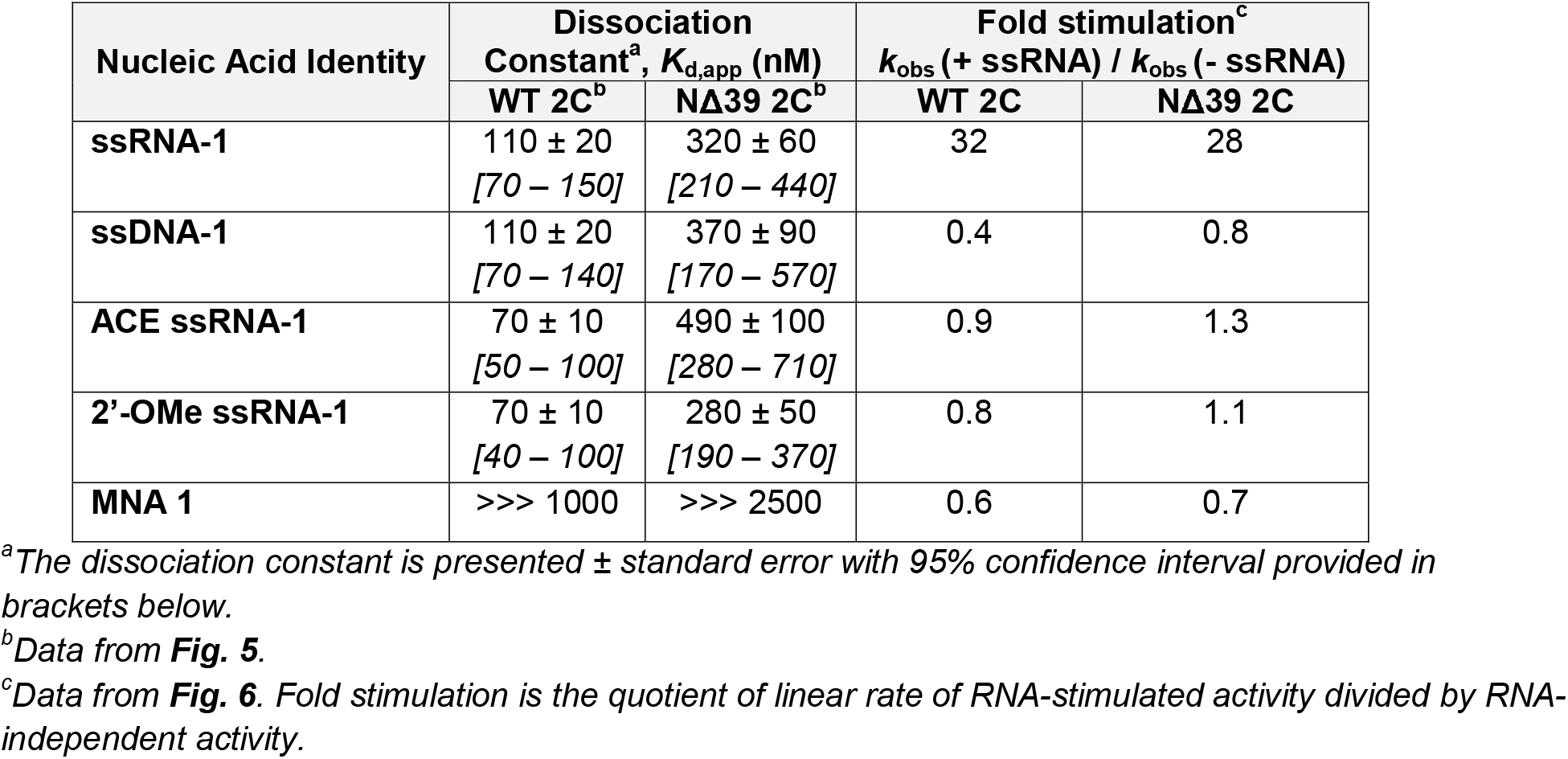
Summary of nucleic-acid binding to 2C and its effect on ATPase activity.

To demonstrate that the site for binding to RNA was the same as that used for binding to DNA, we performed competition experiments. Both unlabeled DNA and RNA competed for binding of labeled RNA with essentially equivalent efficacy for both WT (**Fig. 5D**) and NΔ39 (**Fig. 5E**). Together, these data suggest that a single binding site exists for binding to single-stranded nucleic acid, and that the primary determinant for binding to this site is the phosphodiester backbone.

### PV 2C exhibits RNA-stimulated ATPase activity

We used a thin layer chromatography-based assay to monitor conversion of [α-P]-ATP to [α-P]-ADP as described under Material and Methods. PV WT 2C exhibited low ATPase activity in the absence of RNA but exhibited substantial activity in the presence of a saturating concentration of RNA (**Fig. 6A**) based on experiments shown above in **Fig. 5**. Quantitation revealed a 30-fold increase in ATPase activity of WT 2C in the presence of RNA (**Fig. 6B,** summarized in **Table 2**). We measured the RNA concentration dependence of the activation. In this experiment, WT 2C was present at a concentration of 2 μM 2C (1 μM dimer). We observed a linear increase in ATPase activity when increasing RNA concentrations until saturation, resembling a titration (**Fig. 6C**). This observation is consistent with RNA binding measured by polarization actually reflecting formation of the 2C dimer.

**Figure 6.**
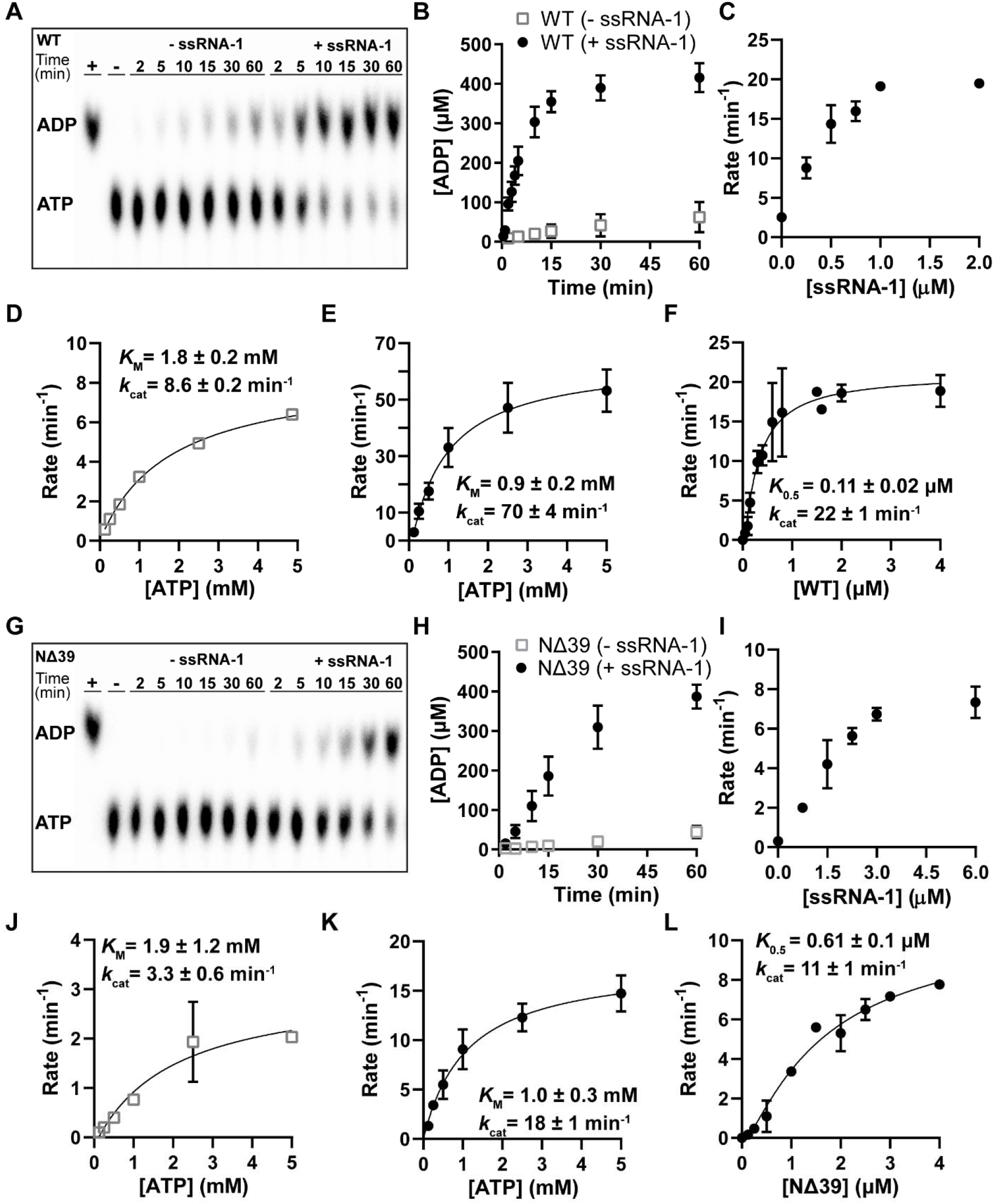
2C exhibits RNA-stimulated ATPase activity. ATPase activity of **(A-F)** WT 2C and **(G-L)** NΔ39 2C. **(A,G)** Phosphorimage of a TLC-PEI plate showing the hydrolysis of [α^32^-P]-ATP to [α^32^-P]-ADP by WT 2C (panel A) and NΔ39 2C (panel G) (2 μM) in the absence or presence of ssRNA-1 (5 μM). HCV NS3 was used as a positive control, denoted (+) and buffer was used a negative control, denoted (-). **(B,H)** Kinetics of ATP-hydrolysis for WT 2C (panel B) and NΔ39 2C (panel H) in the absence or presence of ssRNA-1. Shown are representative time courses using 2C (2 μM), ATP (500 μM), and ssRNA-1 (0 or 5 μM). The observed rates of ATP hydrolysis during the linear phase of the reactions were: WT, 1.4 ± 0.5 μM ADP min^-^ in absence of nucleic acid and 44 ± 3 μM ADP min^-^ in the presence of ssRNA-1; NΔ39, 0.47 ± 0.1 μM ADP min^-^ in absence of nucleic acid and 13 ± 2 μM ADP min^-^ in the presence of ssRNA-1. **(C,I)** RNA concentration dependence of 2C-stimulated ATPase activity for WT 2C (panel C) and NΔ39 2C (panel I). The rates of ATP hydrolysis as a function of RNA concentration were measured and plotted. The resulting data reflect a titration. **(D,E,J,K)** Steady-state kinetic analysis of 2C-ATPase activity was evaluated in the absence or presence of ssRNA-1 (WT, panels D,E; NΔ39, panels J,K). The initial rates of ATP hydrolysis were determined at different concentrations of ATP using 2C (2 μM) and ssRNA-1 (0 and 5 μM). The data were fit to Equation 2 yielding values for *K_m_* and V_max_ reported in each panel. These data are summarized in **Table 3**. **(F,L)** 2C concentration dependence on the steady-state rate of ATP hydrolysis (WT, panel F; NΔ39, panel L). The initial rates of ATP hydrolysis were determined using ATP (500 μM) and a saturating concentration of ssRNA-1 (5 μM). The data were fit to Equation 6 yielding the *K*_0.5_ values reported in the individual panels.

To determine if RNA affects substrate binding and/or the rate-limiting step of the reaction under the conditions used, we performed a steady-state kinetic analysis of ATP hydrolysis in the absence (**Fig. 6D**) and presence (**Fig. 6E**) of RNA. The vast majority of the effect of RNA is on the maximal velocity of the reaction instead of the *K*_M_ for ATP. These data are summarized in **Table 3**.

**Table 3.** Summary of steady-state kinetic constants.

| Substrate | [ssRNA-1]<br>($\mu\text{M}$) | Assay | WT 2C | | NΔ39 2C | |
| --- | --- | --- | --- | --- | --- | --- |
| | | | $K_M$ (mM)<sup>a</sup> | $k_{\text{cat}}$ (min<sup>-1</sup>)<sup>a</sup> | $K_M$ (mM)<sup>a</sup> | $k_{\text{cat}}$ (min<sup>-1</sup>)<sup>a</sup> |
| ATP | 5 | Radioactive<sup>b</sup> | $0.86 \pm 0.2$<br>[0.4 – 1.3] | $70 \pm 4$<br>[63 – 78] | $1.0 \pm 0.3$<br>[0.2 – 1.7] | $18 \pm 1$<br>[16 – 21] |
| ATP | 0 | Radioactive<sup>b</sup> | $1.8 \pm 0.2$<br>[1.4 – 2.1] | $8.6 \pm 0.2$<br>[8.1 – 9.1] | $1.9 \pm 1.2$<br>[-0.6 – 4.5] | $3.2 \pm 0.6$<br>[2.1 – 4.5] |
| 3'-dATP | 5 | Colorimetric<sup>c</sup> | $0.70 \pm 0.1$<br>[0.5 – 0.9] | $27 \pm 0.7$<br>[25 – 29] | $3.5 \pm 0.9$<br>[1.5 – 5.4] | $19 \pm 2$<br>[15 – 24] |

<sup>a</sup>The $K_M$ and $k_{\text{cat}}$ are presented ± standard error with 95% confidence interval provided in brackets below.
<sup>b</sup>Data from **Fig. 8**.
<sup>c</sup>Data from **Fig. 9**.

If the *K*_d,app_ measured using polarization actually reflects formation of the 2C dimer, then under conditions in which RNA is present at a saturating concentration, then we should observe a 2C concentration dependence on the steady-state rate of ATP hydrolysis. In this case, the data should also fit to a quadratic equation with the value of *K*_0.5_ (2C concentration exhibiting half of the maximal stimulation) should be on par with the *K*_d,app_ value measured in **Fig. 5**. Furthermore, the maximal rate should saturate at 2 μM 2C (1 μM dimer) and remain steady until the concentration of RNA is exceeded. The data were consistent with these predictions (**Fig. 6F**). However, the *K*_0.5_ value measured was slightly higher than expected. We conclude that RNA binds productively to a 2C dimer and binding of RNA to a 2C monomer may interfere with formation of a 2C dimer capable of achieving the most active catalytically competent state.

We performed the same series of experiments using PV NΔ39 2C (**Figs. 6G-L**). The conclusions drawn from these experiments were the same as those above for WT. Deletion of the first 39 amino acid residues causes a small but significant reduction in the affinity for ATP and maximal rate of ATP hydrolysis (compare **Figs. 6I,J** to **Figs. 6D,E**). The amount of NΔ39 2C required to observe maximal activity was also higher (**Fig. 6L**), suggesting that the amino terminus may contribute in some way to 2C dimer formation and/or stability.

### A two-step mechanism for RNA binding to PV 2C and activation of its ATPase activity

Because DNA binds to PV 2C as well as RNA, we expected DNA to activate the ATPase activity as well as RNA. This was not observed for either WT or NΔ39 2C proteins (see panels DNA in **Figs. 7A-D**). Interestingly, modification of the 2-hydroxyl also interfered with the ability of such modified RNA to stimulate the ATPase activity of both proteins (see panels ACE and 2’-OMe in **Figs. 7A-D**). Consistent with the inability of MNA to bind to either protein, we did not observe stimulation of the ATPase in the presence of MNA (see panel MNA in **Figs. 7A-D**). The simplest explanation for these observations is that RNA binds to PV 2C using a two-step mechanism. In the first step, binding is driven by the phosphodiester backbone. In the second step, one or more 2’-hydroxyls engage the protein and/or ATP to augment catalysis at the active site.

**Figure 7.**
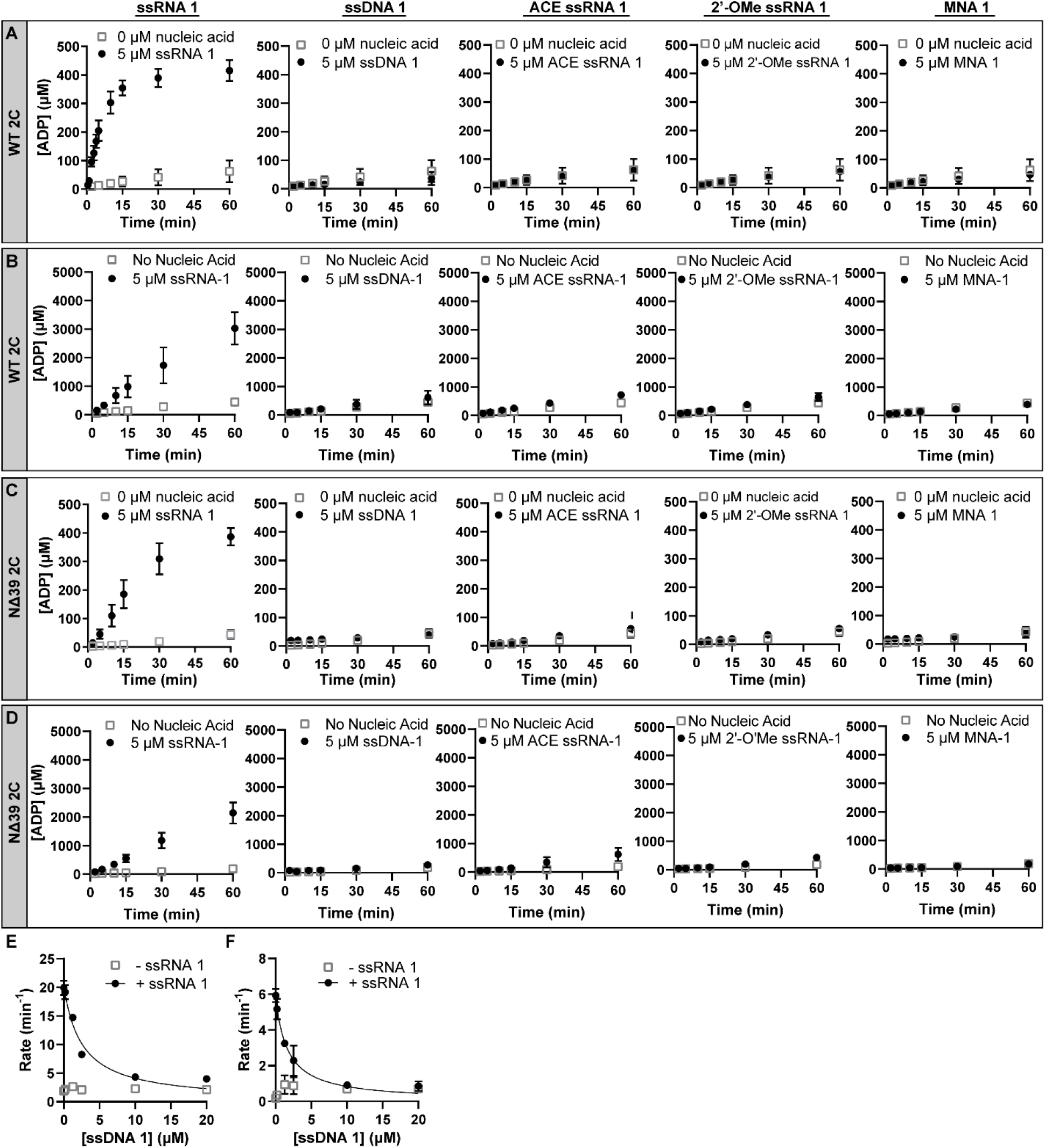
Stimulation of 2C ATPase activity requires a 2’-OH. Kinetics of ATP-hydrolysis for **(A,B)** WT 2C and **(C,D)** NΔ39 2C protein in the absence or presence of nucleic acid (ssRNA; ssDNA; 2’-ACE ssRNA, 2’-OMe ssRNA, and MNA). Representative time courses are shown using 2C (2 μM), ATP (**A,C**: 500 μM; **B,D**: 5000 μM), and nucleic acid (0 or 5 μM). The initial rates of ATP hydrolysis were determined, and the fold stimulation is summarized in **Table 1**. Only the ssRNA-1 stimulated 2C ATPase activity. **(E,F)** RNA-stimulated 2C ATPase activity competition experiments. A fixed concentration of 2C (2 μM; WT, panel C; NΔ39, panel D) was incubated with ssRNA-1 (0 or 5 μM) in the absence (open grey square) or presence (closed circles) of varying concentrations of ssDNA-1. The initial rates of ATP hydrolysis were determined and plotted. The data were fit to Equation 3. Error bars represent the SD (n = 3). Only the RNA-stimulated 2C ATPase activity was reduced with increasing concentrations of ssDNA-1. The stimulation rates and fold-changes presented here are summarized in **Table 2**.

We evaluated the capacity for dsRNA to activate 2C ATPase activity (**Fig. S7**). This experiment used two complementary 19-nt ssRNA sequences (ssRNA-6 and ssRNA-7) with a low probability of forming secondary structure that form a dsRNA (Duplex 3) when annealed (**Fig. S7A**). At 10 μM, each of these ssRNAs only exhibited a two-fold stimulation of the ATPase activity (**Fig. S7B**). Under these same conditions, ssRNA-1 exhibited a 16-fold increase (**Fig. S7B**). These studies suggest that RNA sequence may also govern the affinity of binding to 2C and the concentration dependence of the activation of ATPase activity. Duplex 3 failed to stimulate ATPase activity (**Fig. S7B**), consistent with the inaccessibility of the 2’-hydroxyls for interactions with the protein.

To strengthen the argument that DNA binds to the same site as RNA and that RNA binding to this site activates ATPase activity, we performed competition experiments. DNA outcompeted RNA, reducing the ATPase to basal levels for both WT (**Fig. 7C**) and NΔ39 2C (**Fig. 7D**). This experiment also highlights the fact that the basal activity observed for 2C does not reflect contaminating RNA, because DNA should have competed contaminating RNA away as well (see –ssRNA-1 in **Figs 7E,F**).

### A two-step mechanism for ATP binding to PV 2C

Previous studies of the nucleotide specificity of the enteroviral 2C protein have hinted at the possibility that nucleotides other than ATP may serve as substrates for 2C (17,62). Given the substantial increase in ATPase activity observed in the presence of RNA, we evaluated nucleotide specificity here. Using [α-^32^P]-labeled nucleoside triphosphates, we only detected 2C-dependent formation of the diphosphate product when ATP was used as substrate (**Fig. 8A**). 2’-dATP was not used as a substrate (**Fig. 8A**). To determine if the inability of the other nucleoside triphosphates to serve as substrates was a reflection of their inability to bind 2C, we evaluated their ability to inhibit turnover of ATP. GTP, CTP, and UTP inhibited turnover of ATP (**Figs. 8B-D,G**). Interestingly, the IC_50_ values measured (**Figs. 8B-D,G**) were on par with the *K*_M_ value measured for ATP (**Fig. 6E**).

**Figure 8.**
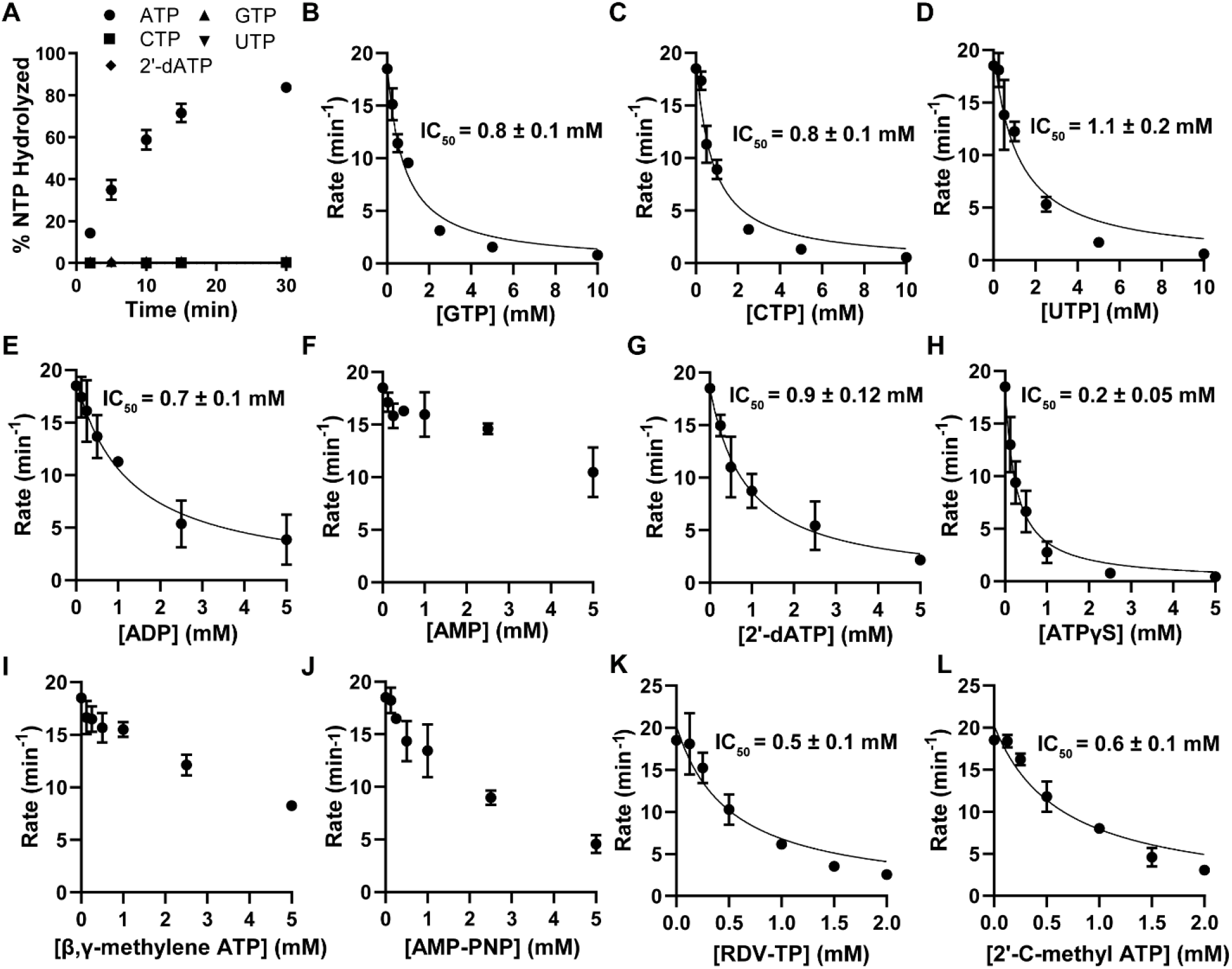
2C-catalyzed ATP hydrolysis occurs via a two-step mechanism. **(A)** Specificity of 2C-nucleoside triphosphatase activity. The kinetics of 2C-catalyzed NTP hydrolysis is depicted using either ATP (•), CTP (■), GTP (▴), UTP (▾), or 2’-dATP (◆) as the substrate. WT 2C (2 μM) was incubated with ssRNA-1 (5 μM) and [α-P]-NTP (1 mM) and the percentage of NTP hydrolyzed was determined and plotted. Error bars represent the SD (n = 3). **(B-J)** 2C ATPase activity competition experiments. A fixed concentration of WT 2C (2 μM) was incubated with ssRNA-1 (5 μM) and [α^32^-P]-ATP (0.5 mM) in the presence of varying concentrations of different unlabeled nucleotides (GTP, CTP, UTP, ADP, AMP, 2’-dATP, ATPγS, β,γ-methylene ATP, or AMP-PNP). The initial rates of ATP hydrolysis were determined and plotted. The data were fit to Equation 3, yielding the IC_50_ values reported in each panel. Error bars represent the SD (n = 3). **(K,L)** Competition of 2C-mediated ATP hydrolysis by the nucleotide analogs, remdesivir-triphosphate (RDV-TP) and 2’-C-methyl ATP. A fixed concentration of WT 2C (2 μM) was incubated with ssRNA-1 (5 μM) and [α-P]-ATP (500 μM) in the presence of varying concentrations of RDV-TP or 2’-C-methyl-ATP. The initial rates of ATP hydrolysis were determined and plotted. The data were fit to Equation 3, yielding the IC_50_ values reported in each panel. Error bars represent the SD (n = 3). *K*_i_ values [] are summarized in **Table 4**.

One explanation for the above observation is that the triphosphate moiety drives the initial binding of a nucleoside triphosphate to 2C. To test this possibility, we evaluated the ability of ADP and AMP to inhibit turnover of ATP by 2C. ADP inhibited turnover of ATP as well as GTP, CTP, and UTP (**Fig. 8E**). AMP was a much weaker inhibitor of ATP turnover (**Fig. 8F**). While ATPγS competed well with ATP, other non-hydrolyzable analogues of ATP did not compete well and may therefore be of limited utility for structural studies (**Figs. 8I,J**). These data are summarized in **Table 4**.

**Table 4.**
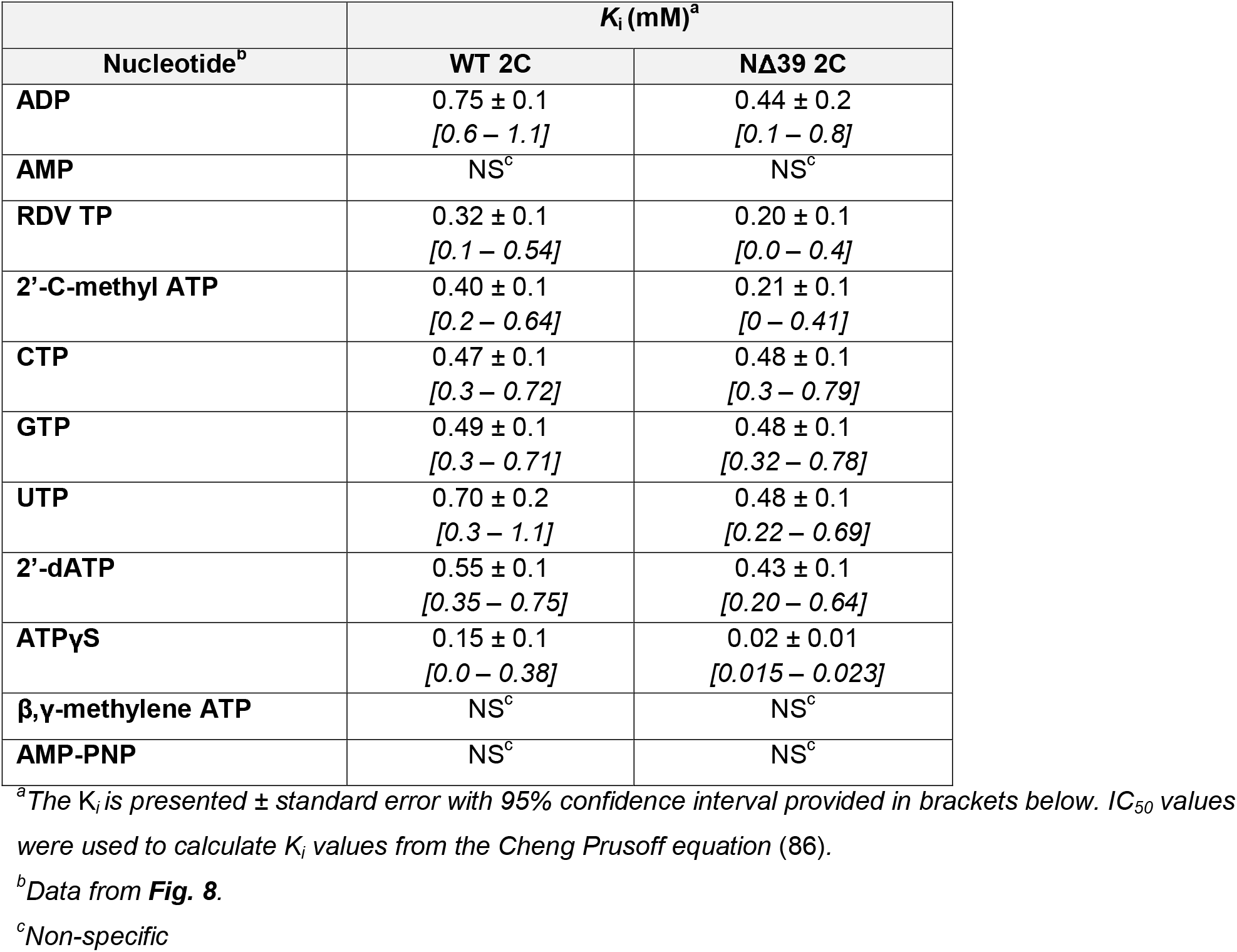
Calculated inhibition constants (*K*_i_) for each nucleotide.

Together, these data suggest that the alpha and beta phosphates are necessary and sufficient for initial binding of nucleotides to 2C. A second step in the binding mechanism would then enable recognition of the base bound and somehow couple the presence of an adenine base to catalytic competence of the active site.

### Determinants of ATP sensed after binding and required for hydrolysis

Our studies make a compelling case for the existence of a conformation of the PV 2C-ATP complex that links binding of the *correct* nucleotide to catalytic competence of the active site. As a first step towards identification of other determinants of ATP interrogated in the second step of binding that are required for catalysis, we have investigated the activity of several analogues of ATP.

First, we evaluated 2’-dATP because a radiolabeled version of this analogue was available. The absence of the 2’-OH prevented hydrolysis to form 2’-dADP (**Fig. 8A**). Other analogues were not available in a radiolabeled form, so we turned to a secondary, colorimetric assay for ATP hydrolysis (**Fig. 9A**). This assay uses a proprietary malachite green-based reagent to detect phosphate directly. The rate of ATP hydrolysis by PV 2C measured using this colorimetric assay was identical to that measured using the radiolabel assay (**Fig. 9B**). The non-hydrolyzable analogues ATPγS and β,γ-methylene ATP were inactive (**Fig. 9D**). Surprisingly, the antiviral nucleotides, remdesivir triphosphate (RDV-TP) (63) and 2’-C-methyl adenosine triphosphate (2’-C-methyl ATP) (64,65) were not substrates (**Figs. 9C,D**), despite both competing well against ATP for binding to 2C (**Figs. 8K,L**). Structures of these analogues are shown in **Fig. 9C**. Not all analogues tested were inactive; 3’-dATP was a good substrate for PV 2C, with a reduced *k*_cat_ value compared to ATP (**Fig. 9E**).

**Figure 9.**
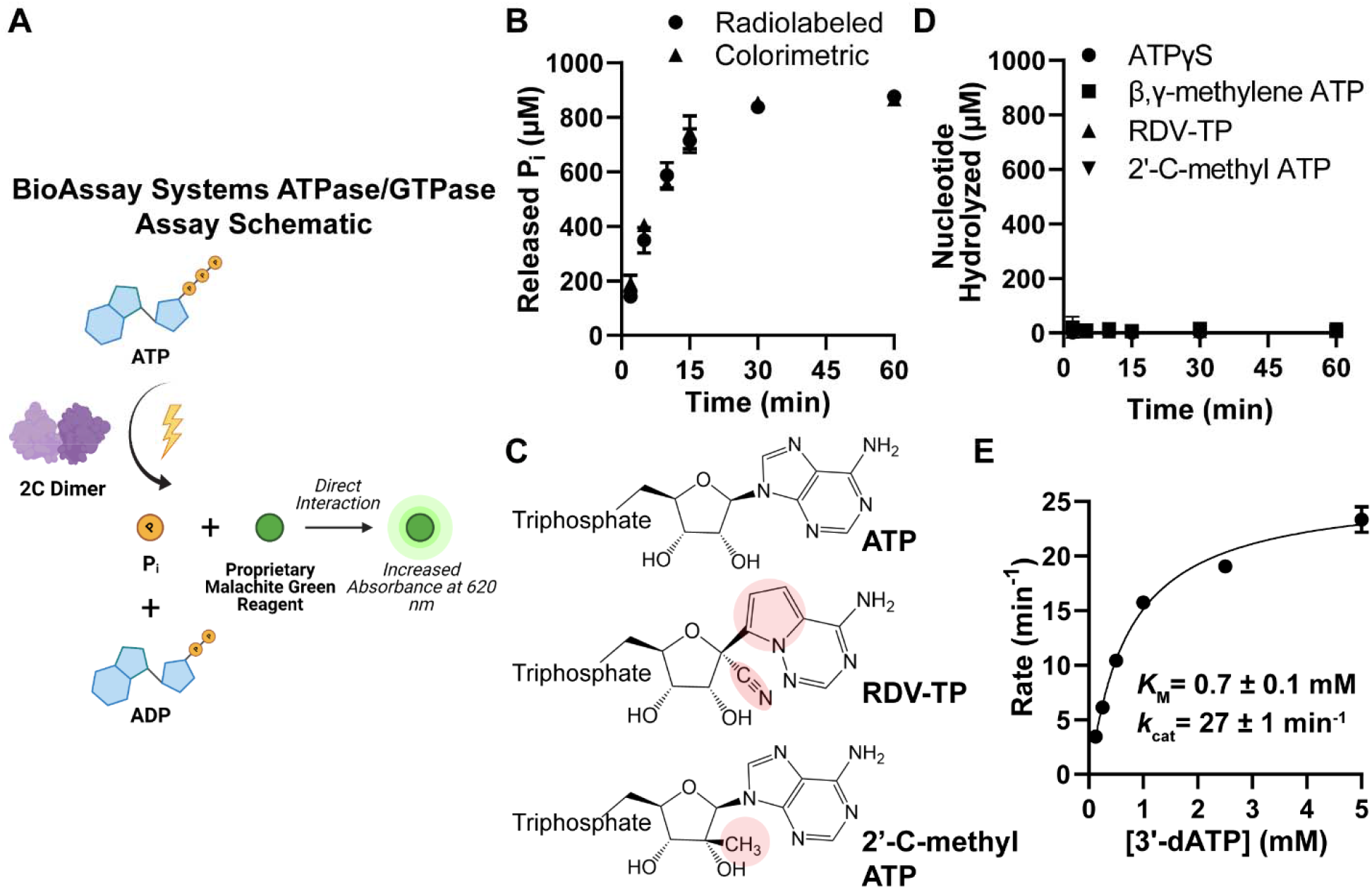
Nucleotide determinants required for 2C-catalyzed hydrolysis. **(A)** Schematic of BioAssay Systems QuantiChrom™ ATPase/GTPase Assay. This assay utilizes a colorimetric method for monitoring ATP hydrolysis. After ATP hydrolysis yielding ADP and phosphate (Pi), a proprietary malachite green compound binds directly to phosphate yielding a green color change that can be detected at 620 nm. The increase in absorbance at 620 nM is directly proportional to the amount of phosphate. Image created with BioRender.com under license. **(B)** Radiolabeled vs Colorimetric ATPase assay. Direct comparison of the kinetics of 2C-catalyzed ATP hydrolysis obtained using radiolabeled ATP or BioAssay Systems colorimetric assay. WT 2C (2 μM) was incubated with ssRNA-1 (5 μM) and ATP (1 mM) and the amount of NTP hydrolyzed was determined and plotted. Error bars represent the SD (n = 3). **(C,D)** 2C does not utilize the adenosine triphosphate analogs, remdesivir-triphosphate (RDV-TP) and 2’-C-methyl-ATP, as substrates. **(C)** Chemical structures for ATP, RDV-TP and 2’-C-methyl-ATP. The chemical moieties that are different from ATP are highlighted. **(D)** Kinetics of 2C-catalyzed NTP hydrolysis. WT 2C (2 μM) was incubated with ssRNA-1 (5 μM) and either ATPγS, β,γ-methylene ATP, RDV-TP, or 2’-C-methyl ATP (1 mM) for up to 60 minutes. The amount of NTP hydrolyzed was determined using BioAssay Systems colorimetric assay. The amount of NTP hydrolyzed is plotted. Error bars represent the SD (n = 3). The molecules ATPγS and β,γ-methylene-ATP was used as a negative control because they are incapable of being hydrolyzed. **(E)** 3’-dATPase activity of 2C. The initial rates of 3’-dATP hydrolysis were determined at different concentrations of 3’-dATP using WT 2C (2 μM) and ssRNA-1 (5 μM) using BioAssay Systems colorimetric assay and is summarized in **Table 3**. The data were fit to Equation 2 yielding values for *K*_m_ and *k*_cat_. Error bars represent the SD (n = 3).

While the structure-activity relationships studied here are far from comprehensive, the relationships enumerated here make it clear that more than the adenine base is essential to formation of a PV 2C-ATP complex competent for hydrolysis.

### PV 2C does not exhibit RNA helicase activity using established helicase substrates

To date, only one laboratory has reported helicase activity for a picornaviral 2C or 2C-like enzyme (26). In the case of the picornaviral 2C, this laboratory suggested that an unidentified, post-translational modification installed by expression of the protein in insect cells bestowed this activity to their protein (26). This conclusion has been challenged by the observation that other picornaviral 2C proteins expressed and purified from insect cells appear to lack a similar post-translational modification and are inactive as helicases (52).

Assaying RNA unwinding generally requires a concentration of dsRNA low enough to prevent reannealing and an enzyme concentration high enough to have all of the dsRNA bound. The experiments described above placed us in a unique position to achieve optimal conditions for unwinding. We used two different unwinding substrates. Duplex 1 has a 9-bp duplex flanked by 20-nt single-stranded tails for loading and translocation of the helicase (**Fig. 10A**). This design permits unwinding regardless of the polarity of the helicase; similar substrates have been used by us in the past (66,67). Duplex 2 is a *forked,* unwinding substrate (**Fig. 10A**). Hexameric helicases have been shown to work best when the displaced strand interacts with the enzyme (67). In these experiments, we used HCV NS3 as a positive control. This enzyme was able to unwind both substrates under two different solution conditions (see panels HCV NS3 in **Figs. 10B,C**). In contrast, PV 2C was unable to unwind either substrate (see panels WT 2C in **Figs. 10B,C**).

**Figure 10.**
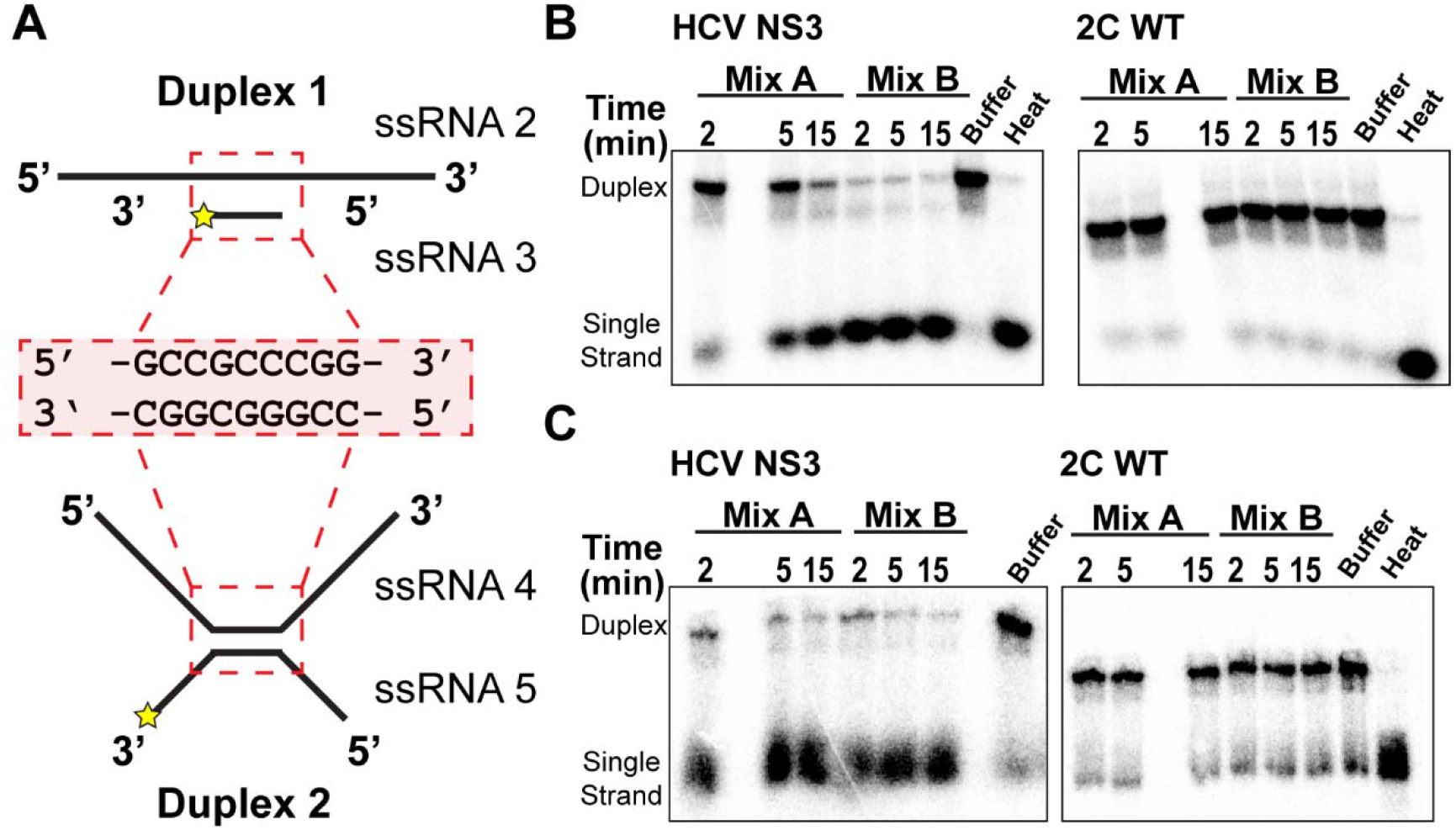
2C does not exhibit RNA helicase activity. **(A)** Schematic of dsRNA Duplex substrates used to assess RNA helicase activity. dsRNA Duplex substrates 1 and 2 are formed using ssRNA 2 & ssRNA 3 and ssRNA 4 & ssRNA 5, respectively. See **Table S1** for RNA sequences. Both Duplex substrates contain a 9-bp region; sequence is highlighted. Duplex 1 is flanked by 20-nt single-stranded tails on the 5’ and 3’-ends of ssRNA 2. Duplex 2 is forked, such that there are two single-stranded tails on either side of the duplex, formed by ssRNA 4 and ssRNA 5. ssRNA 3 and ssRNA 5 were P-labeled, indicated by a star. **(B,C)** RNA helicase activity. RNA unwinding by 2C was assessed by phosphorimaging of native polyacrylamide gels. HCV NS3 (0.5 μM) or WT 2C (2 μM) was incubated with 10 nM Duplex 1 (panel B) or Duplex 2 (panel C) using two different reaction conditions (Mix A and Mix B as detailed under Materials and Methods). The unwinding reactions were quenched after 2, 5, and 15 minutes. HCV NS3 was used as a positive control for RNA unwinding. Buffer indicates quenched reactions without enzyme, and heat indicates reactions heated to 90 °C prior to being quenched, resulting in single-stranded RNA.

### Other enteroviral 2C proteins exhibit RNA-stimulated ATPase activity

A major conceptual advance of the studies reported here for PV 2C is that RNA is required for optimal ATPase activity. To broaden the significance of this study, we have purified the NΔ39 2C derivatives from enterovirus A71 (EV-A71), Coxsackievirus B3 (CVB3), and enterovirus D68 (EV-D68) using a protocol identical to that used for PV as described under Materials and Methods (**Fig. 11A**). All of these enzymes exhibited RNA-stimulated ATPase activity, are not stimulated by ssDNA, and fail to hydrolyze 2’-dATP (**Fig. 11B**).

**Figure 11.**
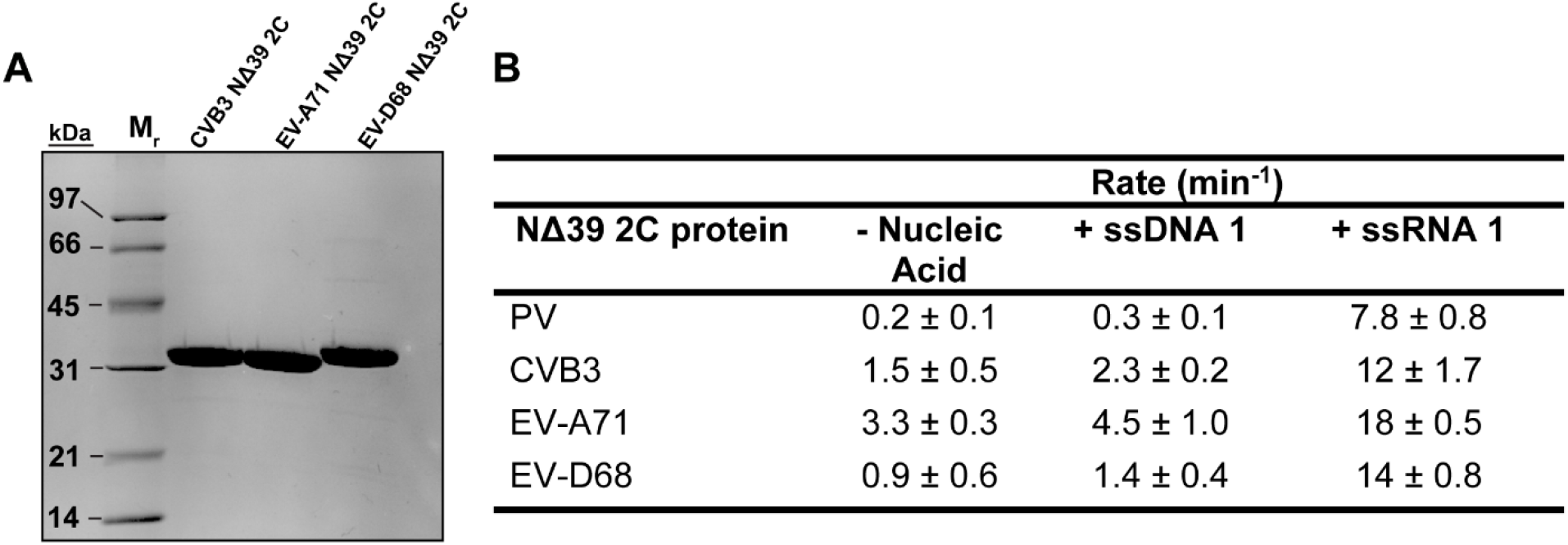
RNA-stimulated ATPase activity observed for members of enteroviral species A-D. **(A)** Purified NΔ39 2C proteins (5 μg each) from Coxsackievirus B3 (CVB3), Enterovirus A71 (EV-A71), and Enterovirus D68 (EV-D68) were visualized on a 15% polyacrylamide gel and was stained with Coomassie. **(B)** RNA-stimulated enteroviral 2C ATPase activity. Rates of ATP hydrolysis were compared between enteroviruses using the indicated NΔ39 2C proteins in the absence or presence of ssRNA. Reactions contained 4 μM 2C, 500 μM ATP, and ssRNA-1 (0 or 10 μM).

## DISCUSSION

Enteroviruses encode several enzymes required for multiplication, the most conserved of which are the 3D gene-encoded RdRp and the 2C gene-encoded AAA+ ATPase and SF3 helicase. A deep understanding of the enteroviral RdRp structure, dynamics, and mechanism exists, although few enteroviral RdRp inhibitors exist (68). In contrast, many enteroviral inhibitors known or suggested to target 2C exist; however, insufficient information exists on 2C biochemistry and biophysics to explain the mechanism of action of any of the known inhibitors. Decades of studies on picornaviral 2C proteins have been published (14,16,17,20). The majority of these have focused on documenting the existence of ATPase activity, demonstrating the use of conserved residues for this activity and virus viability, and/or investigating subunit stoichiometry and organization (13–18,20,23,28,29). Our current mechanistic understanding of the enteroviral 2C ATPase lags far behind viral ATPases and helicases belonging to other superfamilies (41,69–73). The goal of this study was to establish a framework to guide future studies of the specificity, kinetic and chemical mechanisms, and inhibition of the enteroviral 2C ATPase using PV 2C as our model system.

The amino terminus of 2C contains an amphipathic α-helix, the function of which is thought to be attachment to membranes used for genome replication and/or morphogenesis (**Fig. 1D**) (13,74). The hydrophobic nature of this helix is thought to promote aggregation and insolubility. Until this study (**Fig. 1**), full-length 2C protein purified by others was always fused to another protein to permit solubility (17,28,29). One study was able to produce untagged protein by translating an mRNA encoding 2C in the presence of a nanodisc-supported lipid bilayer (75). Another study involved making 2C derivatives that included the full-length protein and soluble portions of the adjacent 2B and 3A proteins (29). Various amino-terminal truncations were made to produce soluble derivatives (27,45,50,51). Deletion of the amino-terminal 33 residues from Foot-and-Mouth-Disease virus (FMDV) yielded an active, well-behaved protein (27). Derivatives from PV and EV-A71 deleted for the first 115 amino acids yielded proteins that could be crystallized, although these proteins were essentially devoid of enzymatic activity (**Fig. S1**). Deletion of the amino-terminal 39 residues of PV 2C and other enteroviral 2C proteins produced active enzymes with overall specificity and activity comparable to the full-length protein (**Figs. 6** and **11**).

Several studies have reported the ability of picornaviral 2C proteins to bind viral and non-viral RNAs, consistent with the expectation that this protein is an RNA helicase (57,58). However, the functional consequence of RNA binding, if any, is not known. Members of helicase superfamily 2, for example the RNA helicase from Hepatitis C virus, binds RNA with high affinity in the absence of ATP but not in the presence of ATP (67). In addition, RNA binding stimulates its ATPase activity (67,73). Toggling between high- and low-affinity states in an ATP-directed manner is thought to couple the directional translocation of the helicase during unwinding to cycles of ATP hydrolysis (67,69,76). Both active and inactive (D177A) forms of PV 2C associated with RNA; however, the presence of ATP had little effect on the interaction with RNA (**Fig. 4**). Binding of RNA to PV 2C and other enteroviral 2C proteins stimulated ATPase activity by more than an order of magnitude (**Figs. 6** and **11**). Steady-state kinetic analysis revealed that the primary effect of RNA is on the maximal rate of the reaction (*k*_cat_) instead of the ATP concentration dependence (compare **Fig. 6D** to **6E** and **6J** to **6K**). At this stage, we do not know the elementary step(s) of the mechanism reflected by *k*_cat_. In some systems, nucleic acid binding stabilizes the closed state of the enzyme, thus promoting catalysis (41,72,77,78). In other non-SF3 systems, the rate-limiting step is release of the phosphate product, which is coupled to the work (translocation, unwinding) performed by the enzyme (67). Regardless of the molecular mechanism, RNA stimulates the activity of 2C to an extent that it may be best to consider 2C as an RNA-dependent ATPase. The fact that others have not reported the RNA dependence may reflect co-purification of RNA and the absence of deliberate assessment of its presence (17,27,28).

AAA+ ATPases and SF3 helicases are usually ring-shaped hexamers or some multiple of hexamers (79). The substrate passes through the central pore in ATP-dependent fashion with processive translocation through the pore facilitated by successive rounds of ATP hydrolysis by one of the six active sites formed at each subunit-subunit interface (79). Most published studies using a purified picornaviral 2C protein have pursued elucidation of its oligomeric structure, most often by using conventional electron microscopy (27–29). However, in no case does it appear that formation of oligomers occurs efficiently

(20,28,29). In our system, the stoichiometry of 2C:RNA was 2:1 (**Fig. 4E**). The value of the *K*_d,app_ for this interaction was in the 100 – 400 nM range (**Figs. 4** and **5**). Because this experiment was performed using polarization, which is sensitive to the mass of the complex, the inability to detect a change in polarization at concentrations as high as 20 μM suggested that the dimer may be a terminal state (**Fig. 4E**). Biophysical experiments, including SEC-MALS (**Fig. 2A**) and SAXS (**Fig. 2C**), were also consistent with formation of only a dimer in solution. Dilution ITC experiments (**Fig. 2B**) were consistent with a dissociation constant in 100 – 400 nM regime. All attempts to use our protein preparations to detect oligomers have only suggested dimers and may require stabilizing complexes with ATPγS as performed in the FMDV system (27). However, even in the presence of ATPγS, the efficiency of oligomerization only approached 15%, and was even less efficient for similar systems using echovirus 30 2C protein (29). We conclude that a functional 2C dimer readily forms in solution and suggest that such a form may have a biological function. If a hexamer is required, then it may need other viral or cellular proteins for assembly (80,81). The empty viral particle represents a possible platform for assembly of a 2C oligomer, reminiscent of the tethering domain observed in SV40 Large T-antigen (**Figs. 3D** and **F**).

Experiments measuring formation of the 2C·RNA complex monitored the 2C-concentration dependence of polarization (**Figs. 4** and **5**). However, it was not clear if the titration measured the concentration dependence of dimer formation followed by RNA binding or sequential binding of 2C monomers to RNA. If formation of the 2C dimer were a prerequisite to binding RNA, then a difference might exist for the observed rate of hydrolysis when the 2C concentration was 10·*K*_d_ compared to that when the RNA concentration was 10·*K*_d_. When the dimer was preformed, we essentially observed stoichiometric activation of the ATPase activity by RNA (**Figs. 6C** and **6I**). However, when RNA was in excess, we observed a normal binding isotherm, with a value of the *K*_0.5_ higher than the equilibrium value of the *K*_d,app_ (**Figs. 6F** and **6L** to **Figs. 5B** and **5C**, respectively). These observations are consistent with a 2C_2_ dimer forming prior to RNA binding. In this case, the affinity of the RNA for the dimer likely has a lower value of the *K*_d,app_ for RNA than the dimer by as much as an order of magnitude. The presence of high concentrations of RNA would yield interactions with monomer, but formation of this complex would interfere with formation of the 2C_2_·RNA complex, resulting in a value of the *K*_0.5_ higher than observed for equilibrium *K*_d,app_ as observed. Loss of the amino-terminal 39 residues increased the *K*_0.5_ value (compare **Fig. 6F** to **6L**), suggesting a role for the amino terminus in formation and/or stability of the 2C_2_ dimer.

Although the 2C dimer bound a variety of nucleic acids (**Fig. 5**) and nucleotides (**Fig. 8**), only RNA activated the ATPase activity (**Fig. 7**) and ATP (**Figs. 8A** and **9**) was the preferred nucleotide substrate. The simplest explanation for this observation is that binding of RNA and ATP to the 2C dimer uses a two-step mechanism. The first step governs ground-state binding and isomerization of this complex yields the activated or catalytically competent state. It is in the second step that *proofreading* of the bound nucleic acids and nucleotide occurs. A similar, two-step mechanism has been shown for binding of RNA to eIF4a family members, DEAD-box helicases contributing to translation initiation (82–84). In the absence of the RNA 2’-hydroxyls, the closed, catalytically competent state cannot be achieved (84). Determinants of ATP specificity are well known for ATPases/helicases (67); however, many helicases will use other nucleotides as well (85). The reason for a strict dependence on ATP is unclear.

The experiments reported herein permit the establishment of the first minimal kinetic mechanism for ATP hydrolysis by the enteroviral 2C protein (**Fig. 12A**). We suggest that the first step is formation of a 2C dimer, 2C_2_, with a *K*_d,app_ in the 0.100 – 1 μM range depending on the disposition of the amino terminus (Step 1 in **Fig. 12A**). Binding of RNA (R) and ATP (T) to 2C_2_ will occur randomly (Steps 2A and 3A or 2B and 3B in **Fig. 12A**) to produce a ternary complex in its ground state, R·2C_2_·T. Formation of this complex is driven by the α- and β-phosphates of the nucleotide, any nucleotide (**Fig. 12B**) and the phosphodiester backbone of nucleic acid, again any nucleic acid (**Fig. 12C**). The ternary complex undergoes a conformational change to produce *R·2C_2_·T* (Step 4 in **Fig. 12A**). ATP undergoes fast hydrolysis in *R·2C_2_·T* and slow hydrolysis in an *activated* binary complex (2C_2_·T* in **Fig. 12A**). We do not believe hydrolysis is occurring in the ground-state complex. If this were the case, then GTP, CTP, and UTP should be substrates. Finally, there are alternative pathways to formation of *R·2C_2_·T* (Steps 3A’-5A’ and 3B’-5B’ in **Fig. 12A**). These are presented for completeness without any evidence of their existence or relevance.

**Figure 12.**
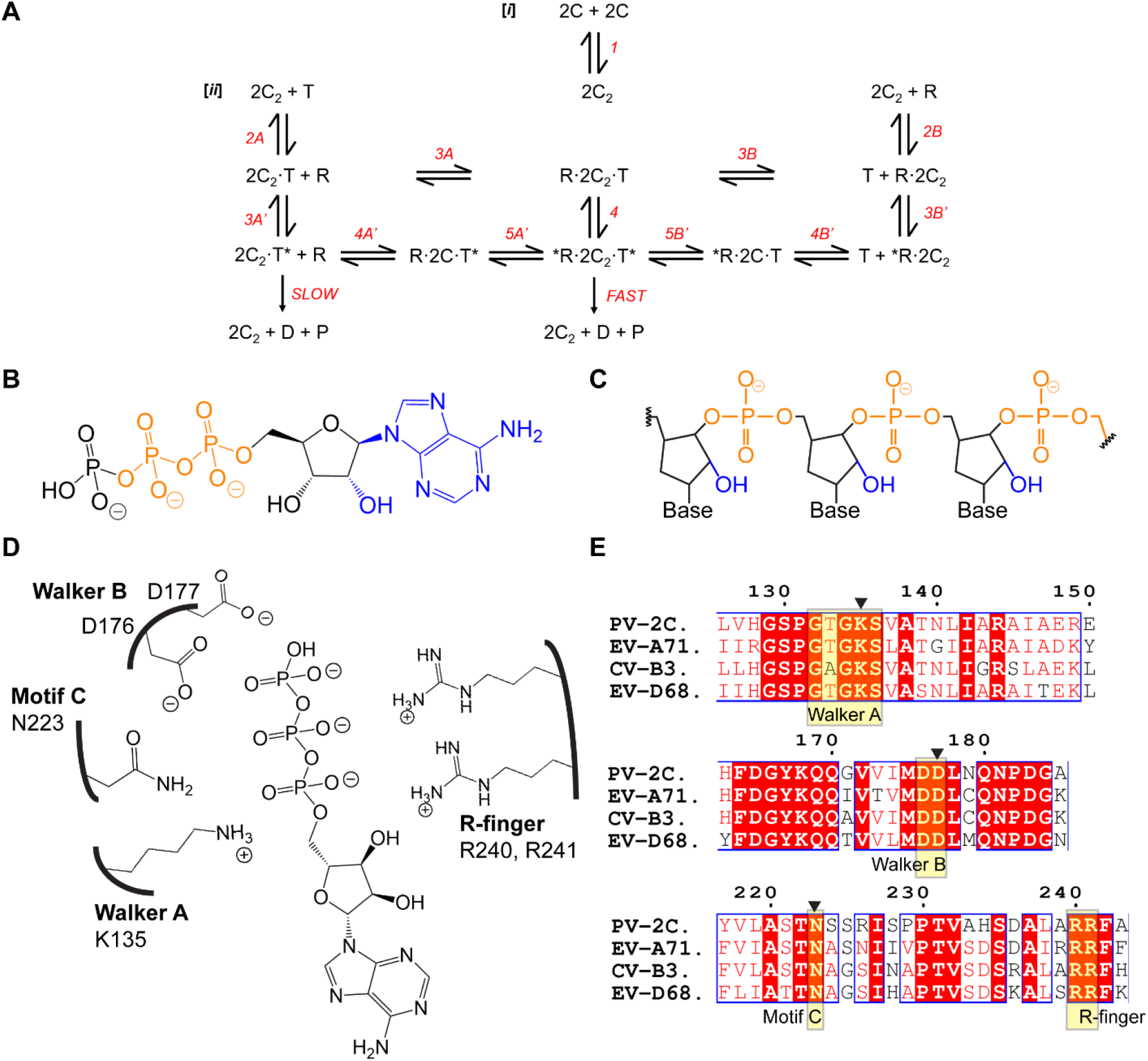
Proposed kinetic mechanism and determinants of specificity and catalytic efficiency for enteroviral 2C ATPase activity. **(A)** Minimal kinetic mechanism for 2C-catalyzed hydrolysis of ATP (T) in the absence and presence of RNA (R) to produce ADP (D) and inorganic phosphate (P). [*i*] Because a dimer is the minimally active form of 2C, experiments were performed under conditions in which the dimer was always present. [*ii*] Binding of ATP and RNA presumably occurs randomly. A major finding here is evidence for a catalytically competent state of the complex (*R·2C_2_·T*) driven by determinants of ATP and RNA that govern ground-state binding (R·2C_2_·T). See text for greater detail. **(B)** Diagram of ATP highlighting the α- and β-phosphates (orange) as primary determinants for binding to 2C and the 2’-OH and adenine base (blue) as determinants sensed after binding to form the catalytically competent state. **(C)** Diagram of RNA highlighting the phosphodiester backbone (orange) as the primary determinant for binding to 2C and the 2’-OH groups (blue) as determinants sensed after binding to form the catalytically competent state. **(D)** Schematic illustrating the proximity of conserved structural motifs to functional groups on the ATP substrates as predicted by other systems (77). The active site is formed at the interface of two subunits. The motifs are: Walker A (K135), Walker B (D177), and motif C (N223), which derive from one monomer of 2C, and the arginine finger (R-finger; R240, R241), which derives from the second monomer of 2C. **(E)** Alignment of sequences comprising the active sites of 2C proteins from PV, EV-A71, CVB3, and EV-D68. Numerous conserved residues exist beyond those with assigned function based on other AAA+ ATPases and SF3 helicases. These conserved residues of the indicated motifs may function to confer the strict specificity of these enzymes for ATP.

Formation and/or stability of *R·2C_2_·T* requires the multiple determinants of ATP: the 2’-OH, the adenine base, among other substituents (**Fig. 12B**). Loss of the 2’-OH or addition of a single methyl group to the 2’-carbon inactivated hydrolysis (**Fig. 9D**). That these observations represent the underpinning of an evolved mechanism is suggested by the fact that the enzyme does not surveil positions of ATP common to all nucleotides; 3’-dATP is a respectable substrate (**Fig. 9E**). Formation and/or stability of *R·2C_2_·T* requires the 2’-OH of RNA (**Fig. 12C**). A structural perspective of this catalytically competent ternary complex does not exist and is difficult to infer. Clearly, the conserved motifs at the active site have the ability to contribute to formation and/or stability of the catalytically competent state (**Figs. 33A** and **12D**). However, these conserved motifs are present in myriad ATPases/helicases, even those that do not exhibit strict specificity for ATP or RNA. Alignment of the enteroviral 2C proteins studied here identify numerous conserved residues beyond the signature motifs that may contribute to the nucleotide and nucleic acid specificity described herein (**Fig. 12E**). We are optimistic that the groundwork established here will facilitate elucidation of high-resolution structures of the catalytically competent state.

The limitations of our study include the use of bacterially expressed proteins and non-viral RNAs. With that caveat noted, whether or not PV 2C or other enteroviral 2C proteins are helicases remains an open question. One study has reported helicase activity for an enteroviral 2C protein (26). The helicase activity is attributed to post-translational modification installed during expression in insect cells (26). This observation has not been reproduced by others (17,28,52). The apparent mass of PV 2C produced in bacteria is equivalent to that produced in PV-infected, human cells (**Fig. S1D**). Substrates for RNA unwinding based on those used by other SF3 helicases (66,67) do not support unwinding by PV 2C (**Fig. 10**). HCV NS3 uses both (**Fig. 10**). If formation of a hexamer is essential for helicase activity, then dimers would be inactive. The translocase activity would be used for encapsidation of the viral genome. As discussed above, perhaps the empty viral capsid would serve as a scaffold for assembly of the hexamer. Indeed, interactions between 2C and the capsid are known and essential for morphogenesis (15,16,18,59). Is there a biological role for a dimeric, RNA-dependent ATPase during the enteroviral lifecycle? Further studies will be required to address this question. If not, perhaps other scaffolds for may exist to assemble hexamers during genome replication. In conclusion, the framework established here will permit elucidation of the specificity and kinetic and chemical mechanisms of the enteroviral 2C protein, as well as enable elucidation of mechanisms of action of inhibitors targeting 2C.

## Data Availability

All data is available upon request.

## Acknowledgements

Our thanks to members of the Cameron Lab for their support. Special thanks to Molly M. Yeager, Ph. D., for edits. Thanks to Dr. Suresh Sharma for his contributions to Fig. S1D. We also thank Ms. Julia Fecko at the X-ray Crystallography Core at the Penn State Huck Institutes of the Life Sciences for contributions to acquisition of data presented in Fig. 2. Finally, we thank Joy Feng for providing remdesivir triphosphate.

