## Supplemental Data for "Enteroviral 2C protein is an RNA-stimulated ATPase and uses a two-step mechanism for binding to RNA and ATP"

### SUPPLEMENTAL FIGURE 1

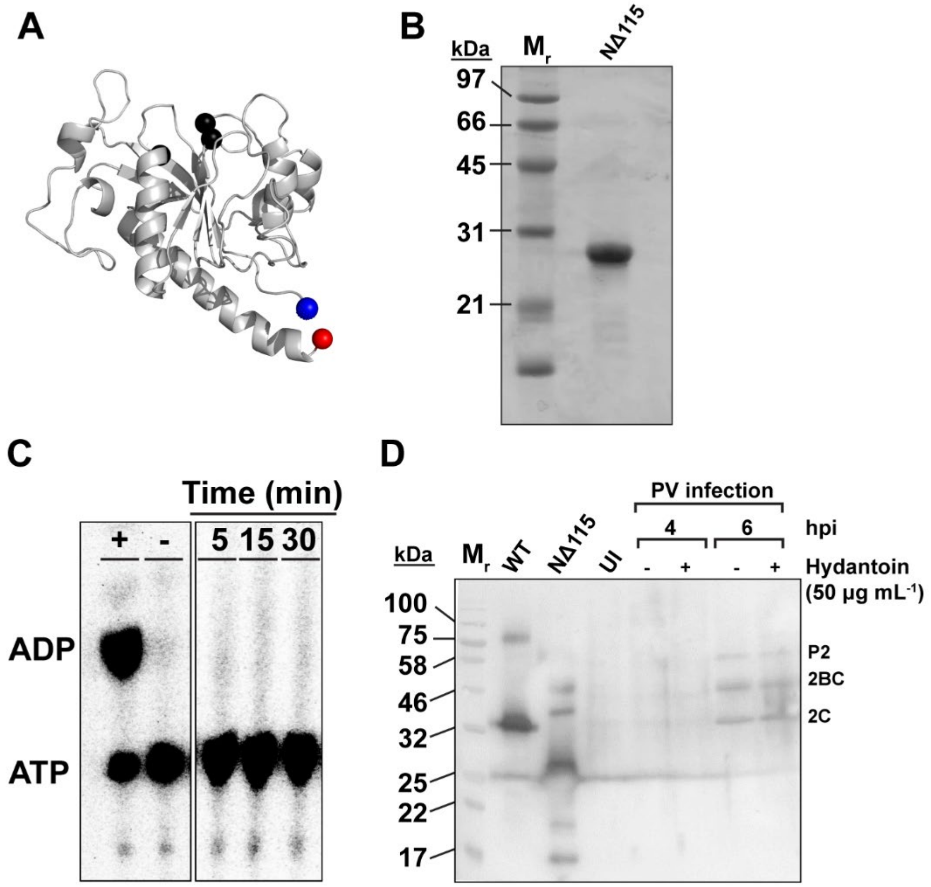


**Figure S1. PV NΔ39 2C: Purified protein and ATPase activity. (A)** Crystal structure of PV 2C in which the amino-terminal 115 residues were deleted (NΔ115) (PDB: 5Z3Q). The amino terminus is shown as a blue sphere; the carboxy terminus is shown as a red sphere; residues required for ATP binding and catalysis are shown as black spheres. **(B)** SDS-PAGE analysis of PV NΔ115 2C protein. Shown is a 15% polyacrylamide gel with 5 µg of purified PV NΔ115 2C protein. Low-range molecular weight markers (Mr) and corresponding molecular weights are indicated. Gel was stained with Coomassie. **(C)** PV NΔ115 2C did not exhibit ATPase activity. Hydrolysis of [$\alpha^{32}$-P]-ATP to [$\alpha^{32}$-P]-ADP by NΔ115 2C was assessed by phosphorimaging of TLC-PEI plates with quenched reaction mixtures. Reactions contained 2C NΔ115 (2 µM), ATP (500 µM), and ssRNA-1 (5 µM). HCV NS3 was used as a positive control, denoted (+); buffer was used a negative control, denoted (-). **(D)** WT PV 2C expressed in bacteria exhibits the same mobility as WT PV 2C produced during infection of HeLa cells. A Western blot was performed using antibodies specific for PV 2C. The mobility of bacterially expressed full-length PV 2C (WT) and NΔ115 2C (NΔ115) was compared to 2C produced during PV infection of HeLa cells. Uninfected cells (UI) are shown next to lysates produced at either 4- or 6 hours post-infection (hpi) in the absence or presence of the assembly inhibitor, hydantoin. Mature 2C (2C) and precursor forms (2BC, P2) were detected at 6 hpi. The mobility of the 2C proteins appear identical.

SUPPLEMENTAL FIGURE 2

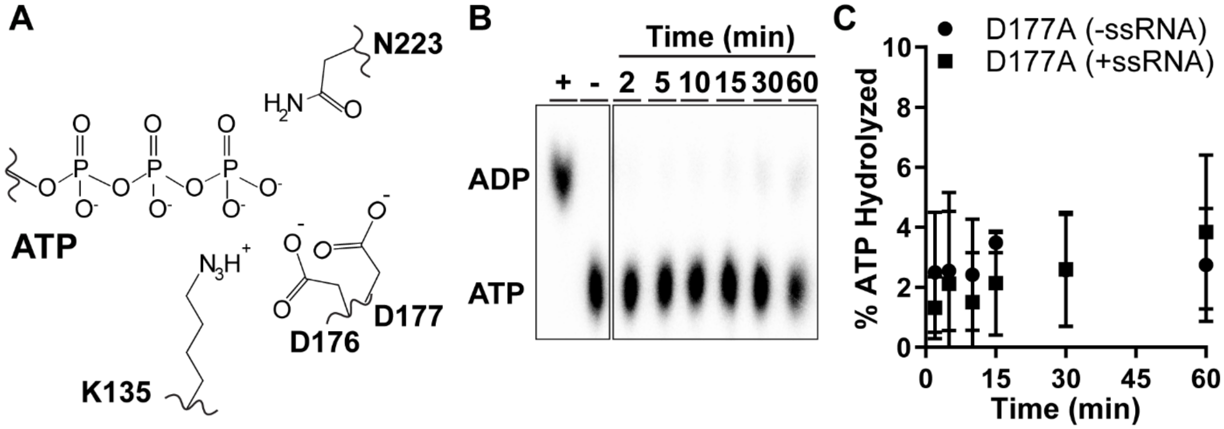


**Figure S2. Genetic inactivation of PV 2C ATPase activity. (A)** Schematic of the residues of PV 2C required for ATP binding and/or hydrolysis. K135 (Walker A motif) contributes to binding of the triphosphate while D177 (Walker B motif) contributes to hydrolysis. Based on other AAA+ proteins, D177 should serve as the general base to activate water for nucleophilic attack of the gamma phosphorous atom. N223 (Motif C) is generally considered the “sensor motif”, and is responsible for detecting the γ-phosphate group of ATP (79). **(B)** PV D177A 2C is debilitated for ATPase activity. Phosphorimage of TLC-PEI plates with quenched reaction mixtures assessing the hydrolysis of [$\alpha^{32}$-P]-ATP to [$\alpha^{32}$-P]-ADP by D177A 2C. Reactions contained D177A 2C (2 μM), ATP (500 μM), and ssRNA-1 (5 μM). HCV NS3 was used as a positive control, denoted (+); buffer was used a negative control, denoted (-). **(C)** Shown are the kinetics of D177A 2C-catalyzed ATP hydrolysis in the absence and presence of ssRNA-1. D177A 2C (2 μM) was incubated with ssRNA-1 (0 or 5 μM) and ATP (500 μM) and the percentage of ATP hydrolyzed was determined and plotted. Error bars represent the SD (n = 3). The hydrolysis rate of ATP with PV 2C D177A in the absence or presence of ssRNA occurred at a rate only slightly higher than background.

**SUPPLEMENTAL FIGURE 3**

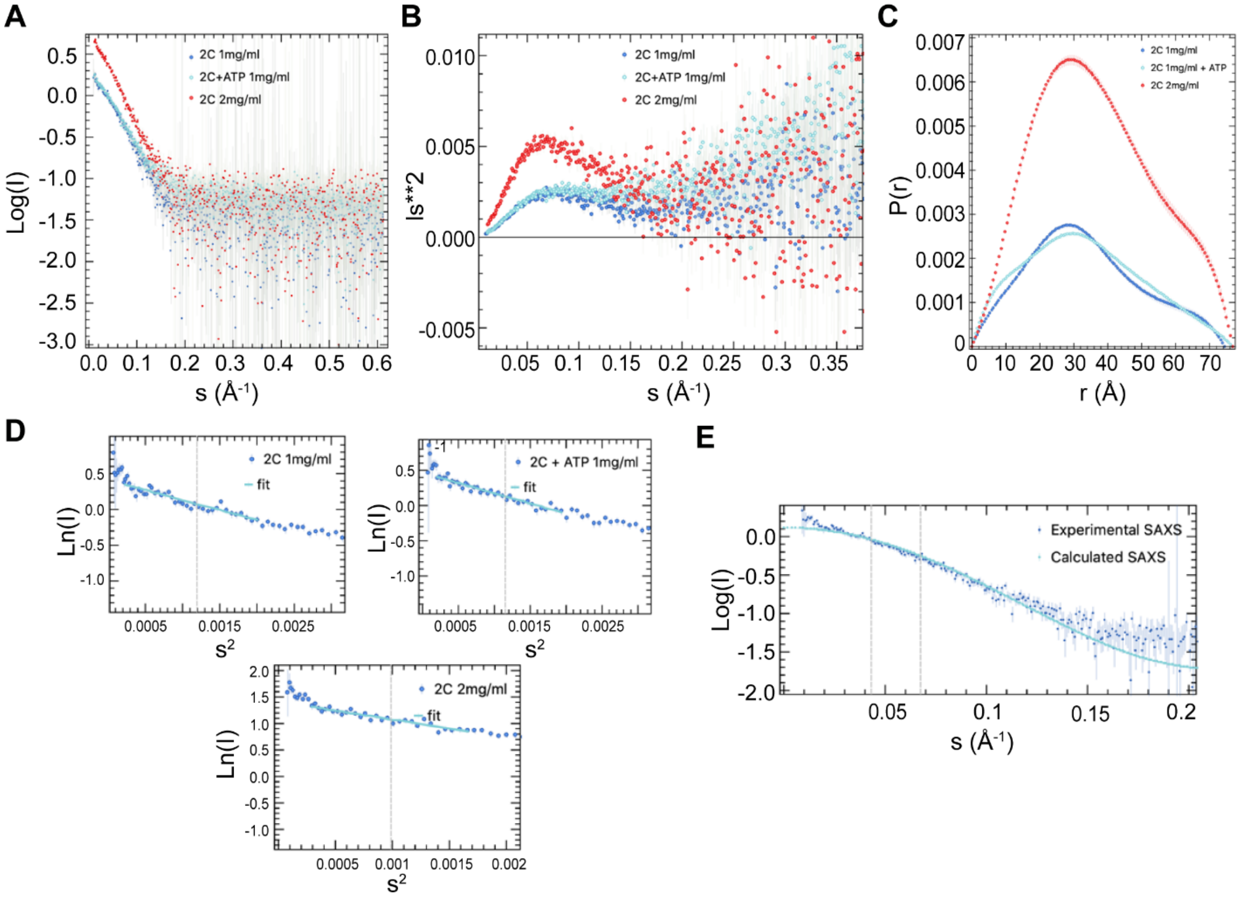


**Figure S3. Analysis of PV 2C by SAXS. (A)** Raw data from small angle x-ray scattering (SAXS) for PV 2C NΔ39 protein at 2 mg ml$^{-1}$ (red) and mg ml$^{-1}$ (blue) in 20 mM HEPES pH 7.5, 5 mM Magnesium Acetate, 1 mM TCEP, 5 % glycerol, and 100 mM NaCl buffer conditions. Data were collected on an in-house Rigaku BioSAXS2000$^{nano}$ for 2C in its apo- and ATP-bound states (cyan, 1:10 molar ratio of 2C:ATP) and analyzed using ATSAS (39). **(B)** Kratky plots derived from SAXS data. (**C)** Pair distance distribution function P(r) overlay. **(D)** Guinear plots for 2C (1 mg ml$^{-1}$) and apo-2C and ATP-bound 2C (2 mg ml$^{-1}$). **(E)** Crysol program overlay of the experimental SAXS profile of 2C (blue; 1 mg ml$^{-1}$) with the calculated SAXS profile as from the predicted dimer model (cyan).

**SUPPLEMENTAL FIGURE 4**

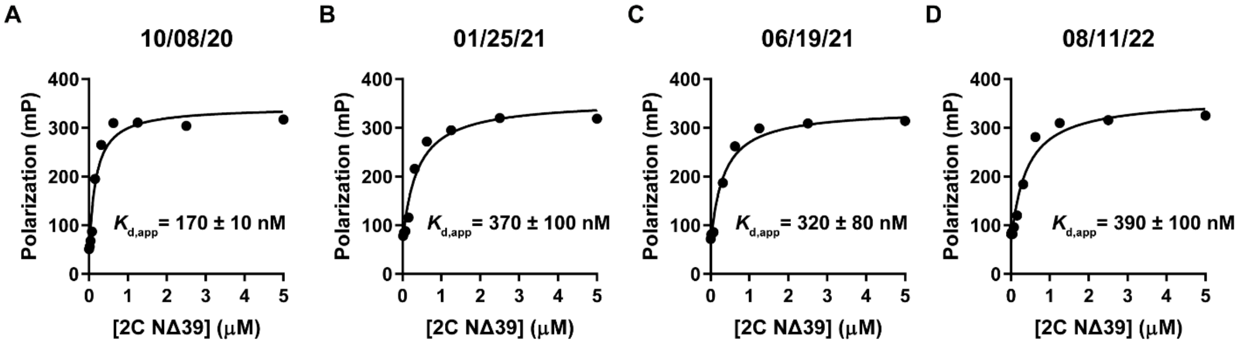


**Figure S4. Fluorescence polarization of multiple biological lots of PV 2C NΔ39.** Representative, individual curves of different biological lots of 2C NΔ39 binding ssRNA-1 (10 nM) from **Fig. 3**. All calculated $K_d$ values were within 3-fold of each other. Lots were purified **(A)** 10/08/20, **(B)** 01/25/21, **(C)** 06/19/21, or **(D)** 08/11/22.

**SUPPLEMENTAL FIGURE 5**

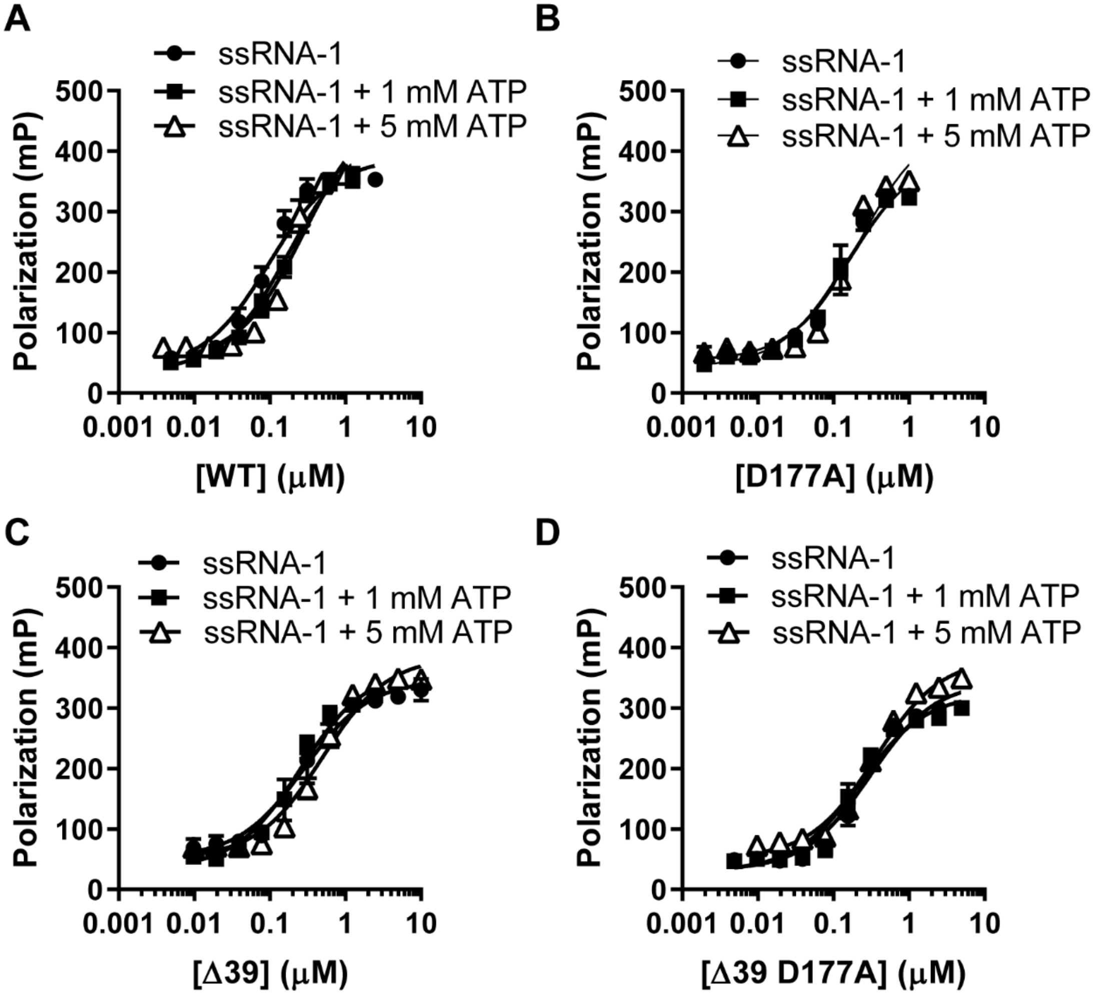


**Figure S5. Logarithmic plots of 2C protein binding to RNA. (A-D)** Data from **Fig. 4** represented as a logarithmic curve and fit using equation 1. 2C protein and the catalytically-inactive D177A 2C bind ssRNA-1 in the absence and presence of ATP (1 and 5 mM). See **Table 1** for $K_{d,app}$ values.

**SUPPLEMENTAL FIGURE 6**

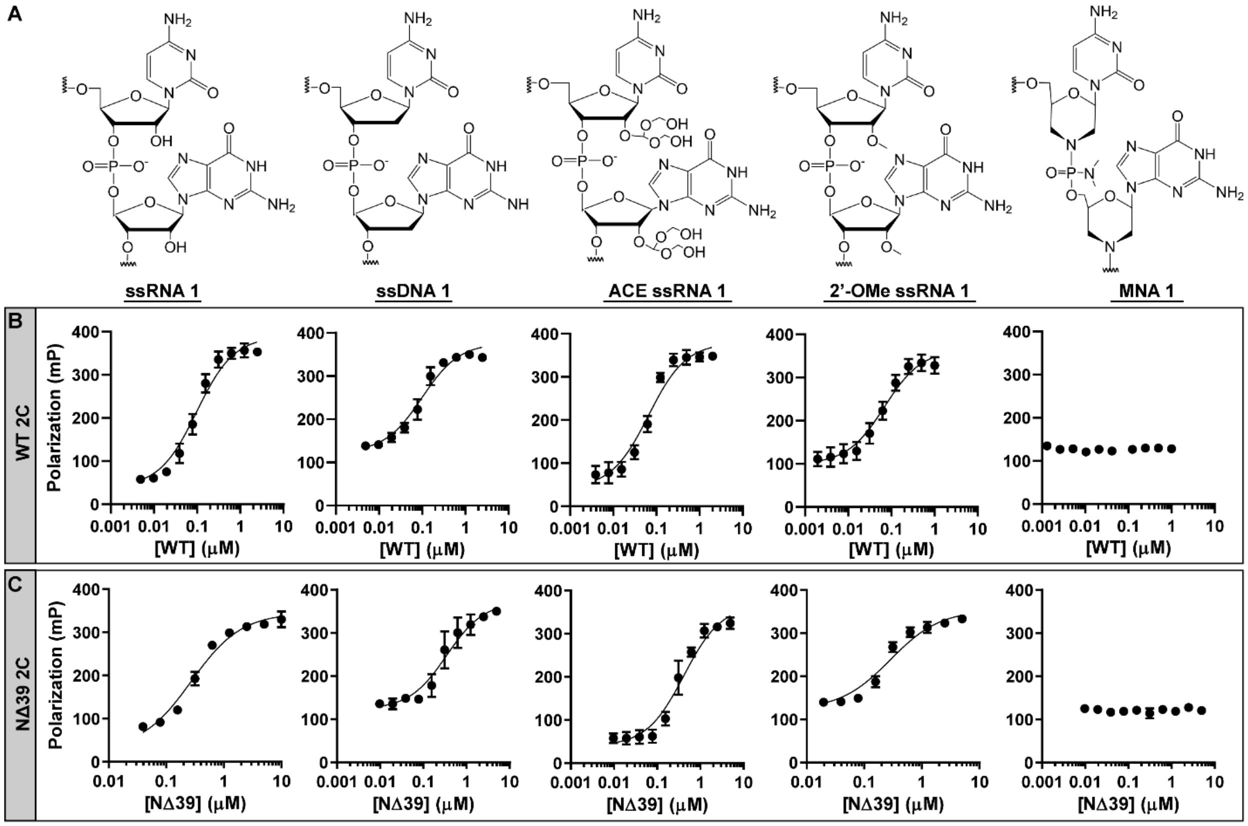


**Figure S6. Logarithmic plots of 2C protein binding to various nucleic acids.** Data from **Fig. 5** represented as a logarithmic curve and fit using equation 1. Represented nucleic acids **(A)** can be bound by **(B)** WT and **(C)** NΔ39 2C.

**SUPPLEMENTAL FIGURE 7**

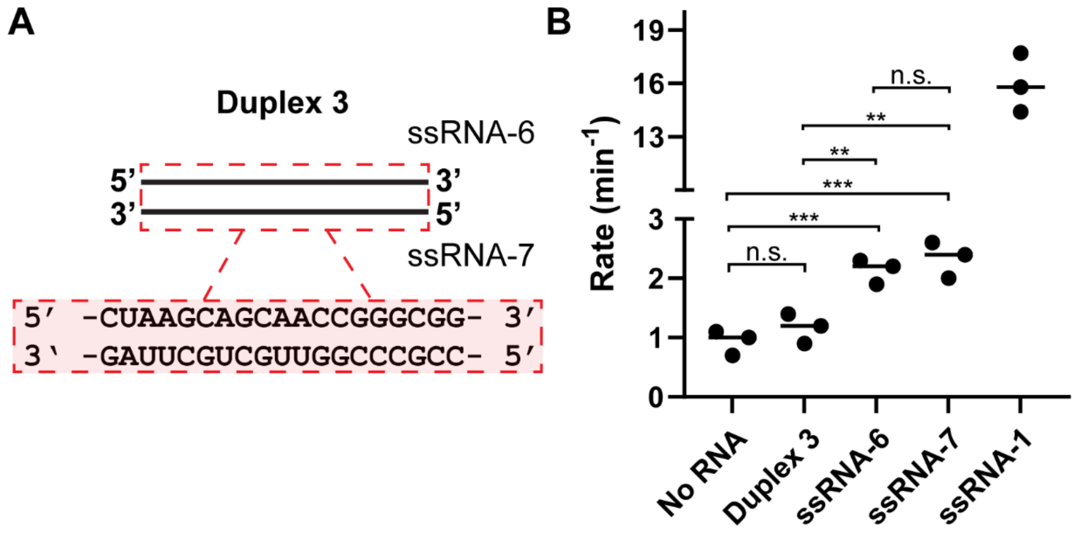


**Figure S7. Double-stranded RNA duplexes do not stimulate 2C ATPase activity. (A)** Because 2C is stimulated by single-stranded RNA, we developed a blunt double-stranded 19mer RNA duplex (Duplex 3) to test for the presence of stimulation via double-stranded RNA. **(B)** RNA-stimulated 2C ATPase activity of the duplex and individual single-stranded RNAs. Rates of ATP hydrolysis were compared between enteroviruses using the indicated NΔ39 2C proteins in the absence or presence of ssRNA. Reactions contained 4 μM 2C, 1 mM ATP, and RNA (0 or 10 μM). For "No RNA" vs "Duplex 3", "No RNA" vs "ssRNA-6", and "No RNA" vs "ssRNA-7", a Student's t-test yielded P-values of 0.2836, 0.0021, and 0.0028, respectively. For "Duplex 3" vs "ssRNA-6", "Duplex 3" vs "ssRNA-7", and "ssRNA-6" vs "ssRNA-7", a Student's t-test yielded P-values of 0.0069, 0.0070, and 0.4018, respectively.

**SUPPLEMENTAL TABLE 1**

**Table S1.** List of nucleic acids used in this study.

| Name | Length (nt) | Nucleobase sequence | Sugar group |
|---|---|---|---|
| **ssRNA-1** | 29 | 5'- GCC-GCC-CGG-CCC-CCC-CCC-CCC-CCC-CCC-CC -3' | Ribose |
| **ssDNA-1** | 29 | 5'- GCC-GCC-CGG-CCC-CCC-CCC-CCC-CCC-CCC-CC -3' | 2'-deoxyribose |
| **ACE ssRNA-1** | 29 | 5'- GCC-GCC-CGG-CCC-CCC-CCC-CCC-CCC-CCC-CC -3' | 2'-orthoester ribose |
| **2'-OMe ssRNA-1** | 29 | 5'- GCC-GCC-CGG-CCC-CCC-CCC-CCC-CCC-CCC-CC -3' | 2'-O-methyl ribose |
| **Morpholino nucleic acid (MNA-1)** | 29 | 5'- GCC-GCC-CGG-CCC-CCC-CCC-CCC-CCC-CCC-CC -3' | Morpholine |
| **ssRNA-2** | 49 | 5'- CCC-CCC-CCC-CCC-CCC-CCC-CCG-CCG-CCC-GGC-CCC-CCC-CCC-CCC-CCC-CCC-C -3' | Ribose |
| **ssRNA-3** | 9 | 5'- CCG-GGC-GGC – 3' | Ribose |
| **ssRNA-4** | 69 | 5'- UUU-UUU-UUU-UUU-UUU-UUU-UU U-UUU-UUU-UUU-GCC-GCC-CGG- UUU-UUU-UUU-UUU-UUU-UUU-UUU-UUU-UUU-UUU -3' | Ribose |
| **ssRNA-5** | 39 | 5'- CCC-CCC-CCC-CCC-CCC-CCG-GGC-GGC-CCC-CCC-CCC-CCC-CCC -3' | Ribose |
| **ssRNA-6** | 19 | 5'- CUA-AGC-AGC-AAC-CGG-GCG-G -3' | Ribose |
| **ssRNA-7** | 19 | 5'- CCG-CCC-GGU-UGC-UGC-UUA-G -3' | Ribose |

**SUPPLEMENTAL TABLE 2**

**Table S2.** List of primers used to produce protein expression plasmids.

| Name | Direction | Nucleotide sequence |
|---|---|---|
| **pSUMO PV 2C WT** | Forward | 5' – GCG-GGA-TCC-GGT-CTC-AAG-GTG-GTG-ACA-GTT-GGT-TGA-AG – 3' |
| | Reverse | 5' – GCG-AAG-CTT-CTC-GAG-TCA-TTA-TTG-AAA-CAA-AGC-CTC-CAT – 3' |
| **pSUMO PV 2C Δ115** | Forward | 5'–GCG-GGA-TCC-GGT-CTC-AAG-GTA-GCA-AAC-ACC-GTA-TTG-AA–3' |
| | Reverse | 5' – GCG-AAG-CTT-CTC-GAG-TCA-TTA-TTG-AAA-CAA-AGC-CTC-CAT-ACG – 3' |
| **pSUMO PV 2C NΔ39** | Forward | 5'-GAA-CAG-ATT-GGA-GGT-GCT-AGA-GAT-AAG-TTG-GAA-TTT-3' |
| | Reverse | 5'–AGT-GGT-GAT-TAT-GGA-CGC-CCT-GAA-TCA-AAA-CCC-AG–3' |
| **pSUMO CVB3 2C NΔ39** | Forward | 5'–GAA-CAG-ATT-GGA-GGT-GTC-AGA-GAA-AAA-CAC-GAG-TTC-CTG-A–3' |
| | Reverse | 5' – CCG-CAA-GCT-TGT-CGA-CTC-ATT-ACT-GGA-ACA-GTG-CCT-CAA-GC – 3' |
| **pSUMO EVA71 2C NΔ39** | Forward | 5'–GAA-CAG-ATT-GGA-GGT-GTC-AAA-GAG-AAA-GTC-GAG-TTT-CTA–3' |
| | Reverse | 5' – CCG-CAA-GCT-TGT-CGA-CTC-ATT-ATT-GGA-AAA-GAG-CTT-CAA-TG – 3' |
| **pSUMO EVD68 2C NΔ39** | Forward | 5'–GAA-CAG-ATT-GGA-GGT-GCT-AGG-GAG-AAA-ATG-AAT-TTG-TGC–3' |
| | Reverse | 5' – CCG-CAA-GCT-TGT-CGA-CTC-ATT-ACT-GAA-ACA-GAG-CTT-CCA-GC – 3' |
| **PV 2C D177A (quikchange)** | Forward | 5'–CTG-GGT-TTT-GAT-TCA-GGG-CGT-CCA-TAA-TCA-CCA-CT–3' |
| | Reverse | 5'–AGT-GGT-GAT-TAT-GGA-CGC-CCT-GAA-TCA-AAA-CCC-AG–3' |
